# Systematics, diversification, and biogeography of Macromiidae (Odonata: Anisoptera)

**DOI:** 10.64898/2026.02.25.708066

**Authors:** Rhema Uche Dike

## Abstract

Macromiidae is a widely distributed lineage of libelluloid dragonflies with a largely allopatric genus-level distribution across the Holarctic, Afrotropical, Australasian, and Indo-Malayan regions. Previous studies involving this family have been complicated by morphological convergence and limited phylogenetic sampling. Here, we present the most densely sampled phylogenetic framework for Macromiidae to date, using Anchored Hybrid Enrichment data from 62 of the 125 described species. Our sampling represents all four genera and major geographic regions, including Libelluloid and Cordulegastrid outgroups. Maximum likelihood recovered three major lineages: *Epophthalmia*, *Phyllomacromia*, and *Macromia* sensu lato, with *Epophthalmia* strongly supported as sister to *Phyllomacromia*. *Didymops* was not recovered as monophyletic and was placed within *Macromia*, although deeper relationships within the *Macromia* complex showed some gene tree discordance. We additionally scored seven male genitalic characters and reconstructed their evolution across a dated phylogeny. We revealed that these traits varied heavily in phylogenetic signal, with some characters supporting the major clades and others showing high degree of homoplasy. Fossil-calibrated divergence time estimation placed the crown origin of Macromiidae in the late Oligocene (24 Ma), with other major intrafamilial divergences concentrated in the Miocene. Historical biogeographic reconstructions consistently supported Afrotropical origins for *Phyllomacromia*, Indo-Malayan centered ancestry for *Epophthalmia*, and a multi-region history for *Macromia* + *Didymops* spanning Indo-Malayan, Australasian, and Nearctic regions. Habitat reconstructions favored lentic ancestry for Macromiidae, and diversification rate variation was best explained by trait-independent models rather than lentic/lotic habitat association.

## INTRODUCTION

The order Odonata, comprising dragonflies and damselflies, has long been a model system for studying evolutionary and biogeographic patterns due to its extensive fossil record, ecological diversity, and high dispersal ability (e.g., Corbet 1999; Goodman et al., 2023; Bybee et al., 2016; Kohli et al., 2021). Macromiidae Needham 1903, a lineage within the superfamily Libelluloidea, is a family of medium to large-bodied dragonflies (∼6 cm to 8 cm in length) with a Holarctic, Afrotropical, Indo-Malayan, and Australasian distribution. The highest diversity in this group is recorded in Southeast Asia and Sub-Saharan Africa (Fig 2). The distribution of the genera within this group shows a largely allopatric pattern. *Macromia* Rambur 1842, is the most widespread genus, and is found in the Nearctic, Palearctic, Indomalayan, and Australasian regions. *Phyllomacromia* Selys 1878, is restricted to the Afrotropical region, including the Seychelles and Madagascar. *Epophthalmia* Burmeister 1839, occurs mainly in the Indomalayan region and parts of Australasia. *Didymops* Rambur 1842, is the most restricted taxon, occuring exclusively in eastern North America.

Historically, Macromiidae was previously considered a subfamily of first Libellulidae (Martin, 1906; Fraser, 1957) and then Corduliidae (Lieftinck, 1971; Kirby, 1890; Orr, 2003) until phylogenetic studies confirmedc it as a distinct family (Carle & Louton, 1994; Ware et al., 2007; Fleck et al., 2008; Michalski, 2012; Bybee et al., 2021).The classification history of Macromiidae was complicated by morphological convergence, given that the first assessments were particularly based on traits such as wing venation (e.g., Fraser 1957; Rehn, 2003); morphological homoplasy is a major challenge in reconstructing phylogenetic relationships. The family designation and name “Macromiidae” was formally proposed by Gloyd (1959) based on diagnoses from three genera: *Macromia* Rambur, 1842, *Didymops* Rambur, 1842, and *Epophthalmia* Burmeister, 1839. She proposed the elevation of what was then called the “Macromia group” to family status.

Currently, Macromiidae consists of four genera (Abbott, 2006-2026; OdonataCentral): *Macromia*, which has been recovered as a sister genus to *Didymops* (Ware et al., 2007; Carle et al., 2015), and *Phyllomacromia*, which has been recovered as a sister genus to *Epophthalmia* (Bybee et al., 2021; Kohli et al., 2021; Kosterin et al., 2025;). *Phyllomacromia* is considered an African genus; a genus-level study by May (1997) examining the status of *Phyllomacromia* and *Macromia* proposed that *Phyllomacromia* should include all African species earlier placed within *Macromia*. *Epopthalmia* is considered an IndoMalayan/Australasian species. An earlier genus-level revision by Lieftinck (1932) revised the status of *Epophthalmia* within what was then the subfamily Corduliinae. Williamson (1909) proposed that North American species previously assigned to *Epophthalmia* should be treated as congenric with *Macromia*, while maintaining *Didymops* as a separate genus despite its close relationship to *Macromia*

An integration of high-throughput sequencing data is needed to resolve the relationships within this group (Kosterin, 2015). Phylogenetic work on Macromiidae has historically been within a broader analyses of Anisoptera or Odonata, with limited taxon sampling within the family. Early multi-gene studies aimed at resolving higher-level relationships consistently recovered Macromiidae as a monophyletic lineage within Libelluloidea, but typically represented it with only a few taxa, failing to test the generic boundaries within the family (Ware et al., 2007; Dumont et al., 2009; Carle et al., 2015). Genome-scale datasets, including anchored hybrid enrichment and transcriptome-based phylogenies, likewise included Macromiidae primarily to stabilize family-level relationships across Odonata (Bybee et al., 2021; Kohli et al., 2021; Goodman et al., 2023; Huang et al., 2025). Targeted molecular studies within Macromiidae have also been restricted to species or regional level. For example, Mehmood et al. (2024) used COI and 16s rRNA data to assess the placement of *Macromia moorei*, recovering it as sister to *Macromia amphigena*, under limited taxon sampling. Recently, a molecular analysis explicitly sampling all four genera, used single mitochondrial and nuclear markers to test the generic limits within Macromiidae. They recovered the family as monophyletic, *Epophthalmia* and *Phyllomacromia* as sister lineages, and *Didymops* nested within *Macromia*, leading to a synonymization proposal (Kosterin et al., 2025).

**Figure 1:**
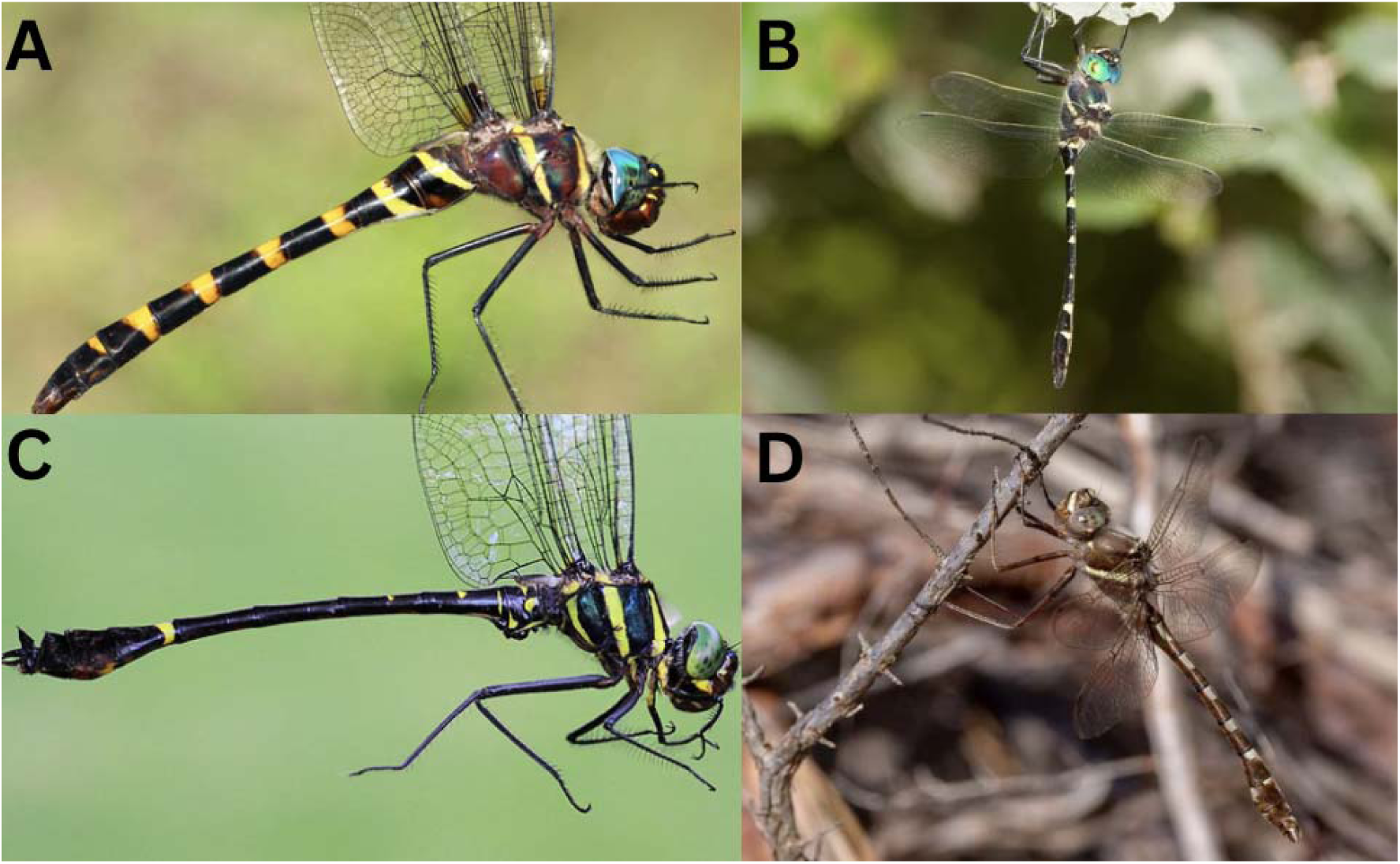
Representative adult individuals from each genus within the family Macromiidae. (A) *Epophthalmia frontalis* (Vietnam; F), (B) *Macromia illinoiensis* (U.S; F), (C) *Phyllomacromia monoceros* (South Africa; M), and (D) *Didymops transversa* (U.S; M). Image credits: A – Tom Kompier; B and D – John Abbott; C – Warwick Tarboton.

Adults in this group can be found patrolling or “cruising” the middle of rivers, lakes, and other water bodies they might inhabit (Novelo-Gutiérrez & Sites, 2024). Despite the common association of most species within this group with rivers and other forms of flowing water, the evolutionary history of habitat use is poorly understood. Indeed, members of Macromiidae are generally associated with lotic habitats, with their nymphs inhabiting clean, well-oxygenated streams and rivers (Corbet, 1999; Needham et al., 2000). However, some species stick to lentic habitats, suggesting potential ecological flexibility that may have influenced diversification within this group. These lentic-lotic transitions may be a driver of evolution by exposing different lineages to distinct selective pressures, especially on nymphal morphology, dispersal behavior, and life history strategies (Moore, 2021; Letsch et al., 2016).

Key morphological synapomorphies for Macromiidae include the sectoral fusion of the arculus for more than half of its length (May, 1997), the extensive contact of the eyes along a majority of their length, with most genera also exhibiting a protrusion on the posterior eye margin (May et al., 2000). In the forewing, the distance between the triangle and arculus is approximately twice that in the hindwing, a diagnostic feature of Macromiidae (May et al., 1997; Garrison et al., 2006). Adult members of this family also share a distinctive sausage-like anal loop with a rounded heel positioned near the base of the hindwing. The anterior hamules are large and erect; posterior hamules not branched and laterally compressed (May, 1997). The nymphal stages of Macromiidae exhibit a suite of unique traits, including broad, flattened bodies with long, spider-like legs (Tennessen, 2019). Nymphs of most genera possess prominent nymphal horns, hand-like tibial spines, and a spoon-like labial mask with deep, jagged teeth on the prementum (Ware et al., 2025).

**Figure 2:**
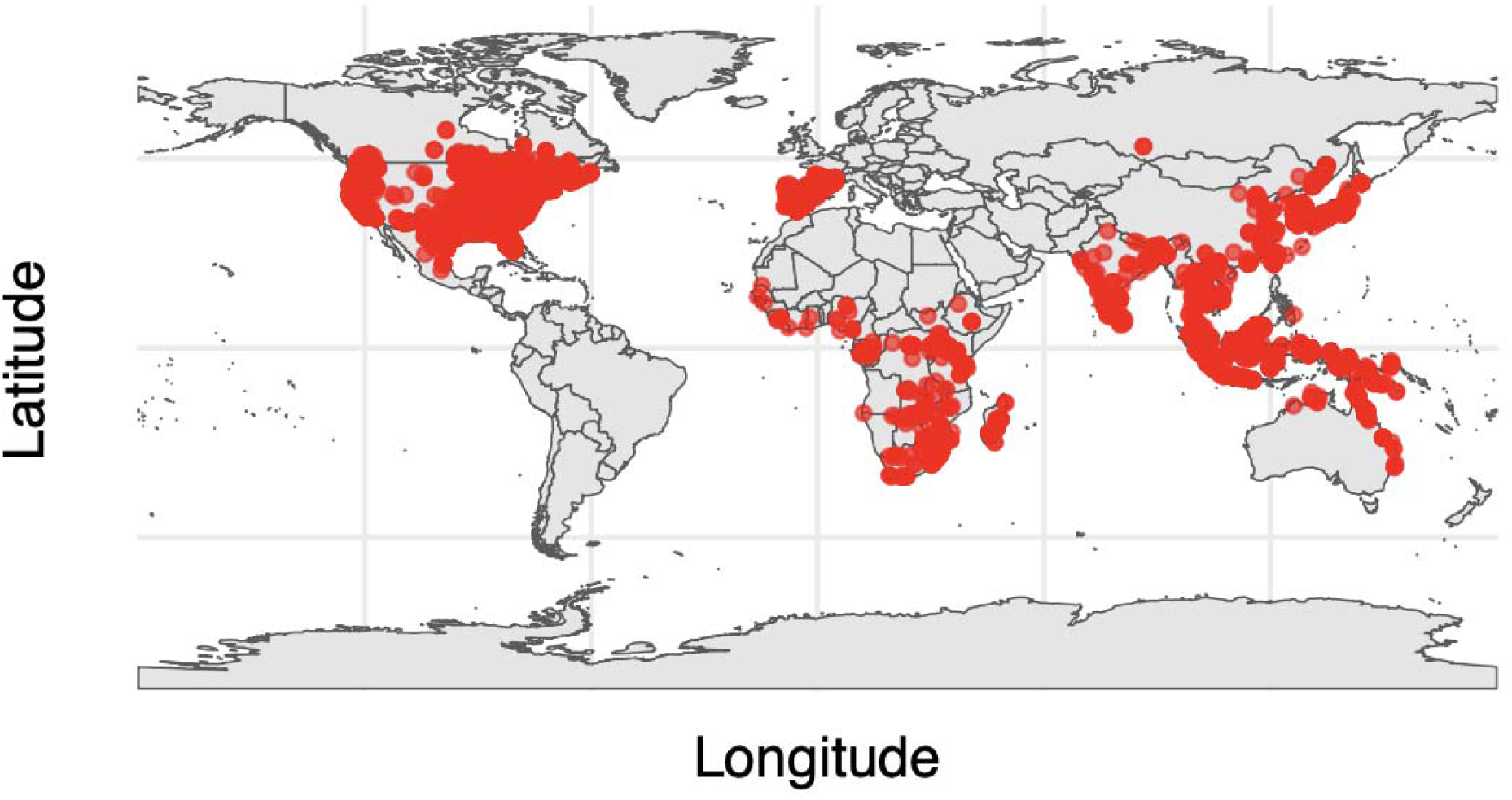
Global occurrence records for Macromiidae. Map showing the geographic distribution of occurrence records for Macromiidae, compiled from the Global Biodiversity Information Facility (GBIF) and iNaturalist databases. Each point represents a verified observation, showing the broad distribution of the family across tropical and temperate regions. The map highlights a pronounced representation in Southeast Asia, sub-Saharan Africa, North America, and Central America.

Advances in molecular systematics have revolutionized our understanding of odonate phylogeny, yet significant gaps remain in resolving genus and species-level relationships, particularly within recently derived lineages such as Macromiidae. Traditional phylogenetic studies based on single orthologous Sanger gene studies have often yielded conflicting topologies due to limitations arising from gene-specific evolutionary rates, horizontal gene transfer, and homoplasy (Delsuc et al., 2005; Jeffroy et al., 2006). Historically, phylogenetic efforts within Macromiidae have been carried out using a few Sanger genes (e.g., Mehmood et al., 2024; Kosterin et al., 2025) while those using subgenomic data have often lacked key taxa (Bybee et al., 2021; Goodman et al., 2023; Kohli et al., 2021;), leading to inconclusive evidence on the relationships within.

With the advent of high-throughput sequencing paired with increased taxon sampling, our ability to obtain higher resolution of deep evolutionary relationships has improved. This provides a more robust framework for understanding diversification processes (Delsuc et al., 2005). This resolution also enables hypothesis testing for trait evolution and conservation of habitat association over time. More specifically, this advent introduces a more reliable backbone to uncover biogeographic patterns within Macromiidae, which have remained largely unexplored, despite their cosmopolitan distribution and ecological diversity. Given the strong habitat specialization of this group to lotic environments in most lineages, it presents an excellent model to investigate phylogenetic niche conservatism, testing whether ancestral habitat preferences have constrained or facilitated diversification over time.

Here, using a targeted enrichment approach, we: (1) carry out a phylogenetic reconstruction of Macromiidae using a combination of high-throughput and morphological data, (2) reveal the historical biogeographic processes shaping the current distribution of the family, (3) estimate divergence times to provide insights into the temporal framework of diversification events within this lineage, (4) reconstruct the ancestral state of habitat preference (lentic/lotic), providing insight into the ecological transitions that may have accompanied lineage diversification within the group. This study provides a comprehensive phylogenomic investigation of Macromiidae, utilizing high-throughput sequencing data, morphological characters, and ecological data to refine interspecific relationships and explore the macroevolutionary trends influencing their evolution.

## MATERIALS AND METHODS

### Taxon Sampling

Specimens were acquired from the Florida State Collection of Arthropods (FSCA), American Museum of Natural History (AMNH), Naturalis Museum of Natural History (NMNH), and Brigham Young University (BYU). Our current analyses include sequences of 62 of 125 species, including all 4 genera included in the family (Supplemental table 1), and this boasts the most robust taxa sampling in this group to date. We selected outgroups from other members of the superfamily Libelluloidea (Libellulidae, Synthemistidae, Corduliidae) and Cordulegastridae (*Cordulegaster*), representing the entire superfamily Cavilabiata.

### DNA Extraction, Probe Design, Sequencing

Legs were pulled from the specimens, and genomic DNA was extracted using a Zymo Mini-prep kit following the manufacturer’s protocols, and the DNA yield was quantified using a Qubit 4 Fluorometer. Extracts were shipped to RAPID Genomics (Gainesville, Florida) for library preparation and sequencing using custom-made Anchored Hybrid Enrichment probes modified and detailed in Bybee et al. (2021). The probes designed contained 1,306 loci, including AHE and functional loci (Bybee et al., 2021; Goodman et al., 2023). These probe sets were created using 941 exons shared across Insecta from 24 odonate transcriptomes and 2 assembled genomes from Bybee et al. (2021) (Goodman et al., 2025). To achieve robust coverage and a solid backbone, we sequenced at least one representative from each genus using the full probe set (500kb), and for the remaining taxa, we sequenced using a subset of 92 loci (20kb). The raw reads used in this study can be accessed via the Dryad digital repository: xxxxx and the information on locus coverage:

### AHE Data Processing

We obtained the raw reads, trimmed adapters using *fastp* (Tang & Wong, 2001), and confirmed the quality of the reads using *multiQC* (Ewels et al., 2016). Each individual locus was assembled using an iterative baited assembly method with *SPAdes* (Prjibelski et al., 2020) and a reference genome assembly of *Tanypteryx hageni* (Anisoptera: Odonata) (Tolman et al., 2023), then screened each locus for orthology by ensuring that the locus did not have BLAST hits to multiple places in the genome, and ensuring the best BLAST hits between the reference and the query sequence (Breinholt et al., 2018). To ensure accurate homology inference and phylogenetic accuracy, we removed taxa that recovered less than 4 loci. Given the potential for noise and failed alignments (Goodman et al., 2024), our analyses included the probe region only.

### Phylogenetic Analyses and Dating

We generated a multiple sequence alignment for the probe and flanking regions and aligned them using MAFFT in *MAFFT* v7.475 (Katoh & Standley, 2013). We concatenated the alignments into a supermatrix using FASconCAT (Ku□ck & Meusemann, 2010), which resulted in 94,657 parsimony-informative sites. An initial partitioning scheme was generated using relaxed clustering in IQ-TREE v2.1.3 with the model fixed to GTR□+□G for each subset, and the best partitioning scheme and models of nucleotide substitution were then refined using *PartitionFinder2* and *ModelFinder* (Lanfear et al., 2017; Kalyaanamoorthy et al., 2017). We concatenated the alignments into a supermatrix using *FASconCAT* (Ku□ck & Meusemann, 2010), which had 37,392 parsimony-informative sites. Using a relaxed clustering model in *Partitionfinder2* and *ModelFinder*, we estimated the best partitioning scheme and models of nucleotide substitution (Kalyaanamoorthy et al., 2017), and reconstructed a maximum likelihood (ML) tree with 1,000 bootstrap replicates using *IQ-TREE* v2.0 (Minh et al., 2020). We selected our outgroups based on sister lineages within the superfamily Libelluloidea, helping to highlight the relative position of Macromiidae.

Given the number of loci used in this study, we presumed there might be cases of gene tree discordance and potentially incomplete lineage sorting (ILS). To assess this genealogical concordance in our dataset between the gene trees and the species tree, we carried out a gene concordance factor (gCF) and site concordance factor (sCF) analysis in *IQ-TREE* v2.0 (Minh et al., 2020). gCF quantifies the proportion of genes/gene trees that support a particular branch in the species tree, providing insight into tree conflicts due to processes like incomplete lineage sorting. sCF, on the other hand, measures the proportion of alignment sites that support a branch, providing information on phylogenetic signal consistency. These analyses helped to uncover the underlying discordance within the data that might lead to topological variation.

We estimated divergence times using the approximate likelihood method implemented in MCMCTree (*PAML* v4.9j; Yang, 2007). Divergence dating was performed on the maximum likelihood phylogeny inferred from our dataset, using a fixed topology in MCMCTree. Our fossil calibration selections followed best practices recommended by Ksepka et al. (2015) and Parham et al. (2012), incorporating fossil information from the Paleobiology Database and relevant existing literature (Goodman et al., 2025; Kohli et al., 2021, Table 1). The root age was constrained with soft bounds of 89.5-158.1 Ma, using 89.5 Ma as a minimum bound for the Libellulidae+Corduliidae lineage and 158.1 Ma as a conservative maximum corresponding to earliest Cavilabiata fossils (Huang and Nel, 2007). We applied additional internal soft bounds at three nodes; the *Epophthalmia*+*Phyllomacromia* clade (15.98-20.45 Ma), the *Macromia*+*Didymops* clade, an internal split within *Epophthalmia* (13.82-15.98 Ma), and an internal split within *Macromia* (0.129-0.774 Ma). additional internal calibrations including Libellulidae*+*Corduliidae at 89.5 Ma (Fleck et al., 1999) and node-specific calibrations for *Macromia pilifera* (Lin, 1982), *Epophthalmia zotheca* (Zhang, 1989), and *Epophthalmia biordinata* (Lewis, 1969). Analyses employed the HKY85 substitution model with five discrete gamma categories (ncatG = 5). Among-lineage rate variation was modeled with an independent-rates relaxed clock (clock = 2). Gamma priors were assigned to substitution model parameters (kappa_gamma = 6,2; alpha_gamma = 1,1), the overall substitution rate (rgene = 2,2), and the variance of rate drift (sigma2_gamma = 1,10). Our birth-death process priors assumed equal speciation and extinction, with the sampling proportion set at 49.2% (λ = 1, μ = 1, ρ = 0.492). Posterior estimates were obtained from four independent MCMC chains, each run for 500,000 samples with a burn-in of 10,000 iterations, sampling every five cycles. Fine-tune settings were adjusted to optimize acceptance rates across node times, substitution rates, and variance parameters. We assessed convergence using Tracer v1.7.2 (Rambaut et al., 2018) by inspecting trace plots and ensuring effective sample sizes (ESS > 200) across all parameters. Consistency among the independent runs confirmed adequate mixing, and posterior divergence times were summarized as means with 95% highest posterior density (HPD) intervals. Node ages were therefore calculated as posterior mean divergence times (in Ma) with 95% HPD intervals after burn-in.

**Table 1:**
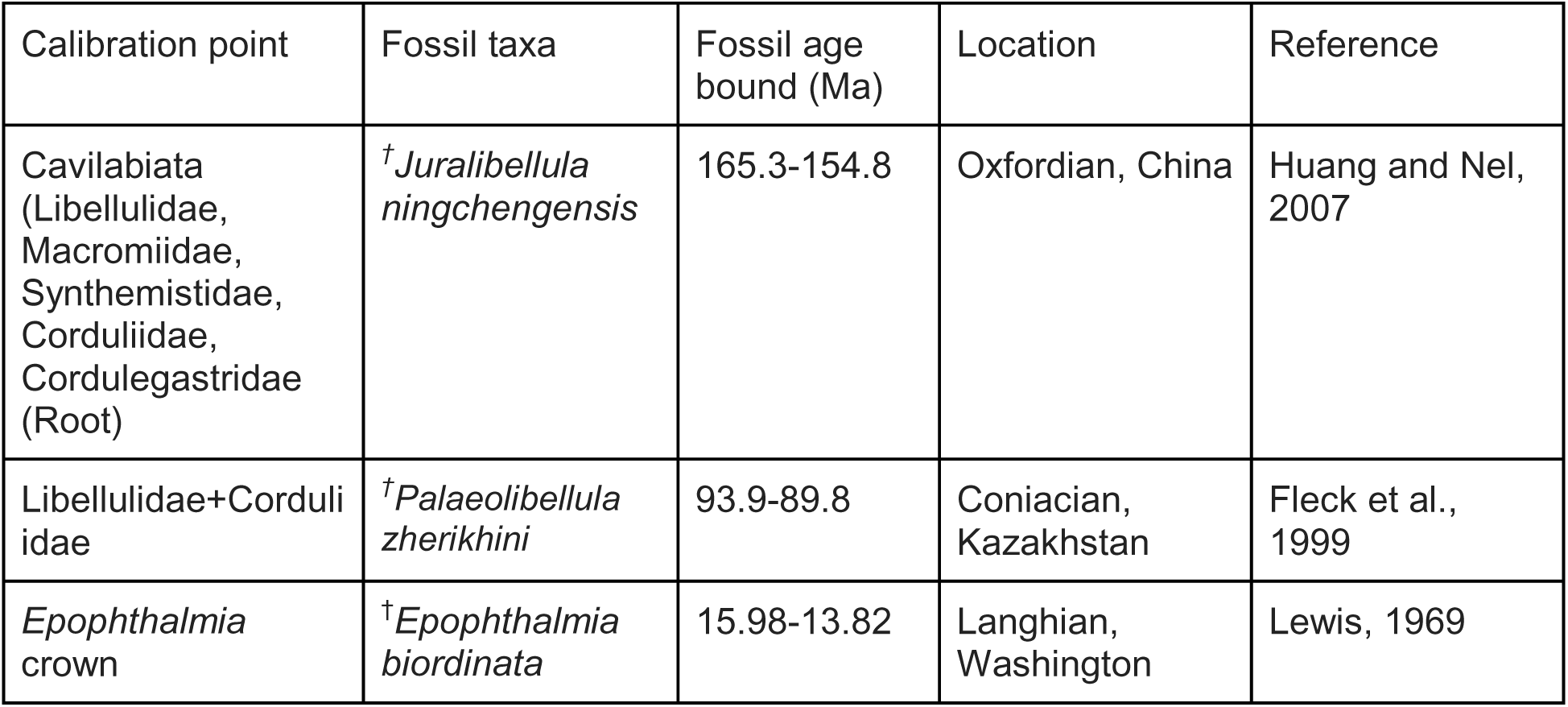
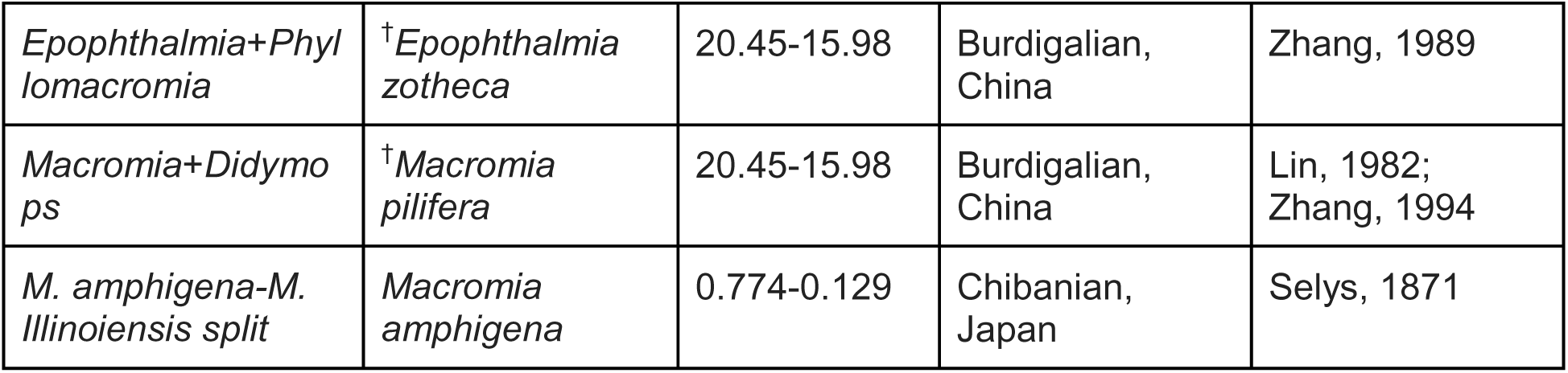
Fossil calibration points used for divergence time estimation. Ages are reported as stratigraphic ranges in million years ago (Ma), with bounds treated as soft minimum and maximum constraints. Taxonomic and stratigraphic justifications follow published diagnoses and original descriptions. Extinct taxa are marked with †

### Morphological assessment

Our morphology matrix consisted of seven male genitalic traits from May et al. (1995) and an unpublished matrix by the same author, for which we verified all state assignments using 34 SEM images. The characters included genital lobe shape, genital ligula shape, penis flagella number, presence of a penis basal lobe, size of penis segments 3 and 4, epiproct length, and epiproct tip width (Supplemental table 2). These structures have been hypothesized to carry phylogenetic signal due to their highly species-specific nature and rapid evolution (Williams et al., 2024; Song & Bucheli, 2010). These characters were coded as unordered, discrete multistate traits. Polymorphic observations were treated as ambiguous states (0&1), and missing data were coded as “?”.

To quantify the phylogenetic signal and degree of homoplasy for each trait, we calculated the Consistency Index (CI) and Retention Index (RI) using the R package phangorn (Schliep, 2011). RI values range from 0 (fully homoplastic) to 1 (fully synapomorphic), CI values range from 0 (high homoplasy) to 1 (perfect consistency of traits).

For each trait, using the *ace* function in ape, ancestral states were reconstructed across the dated phylogeny using ML under the equal-rates (ER) model. Reconstruction outputs were visualized with proportional likelihood pies at nodes. Node certainty was categorized by the ML value: high (≥0.8), moderate (0.6-0.79), and low (<0.6). Observed tip states were shown as filled circles and missing data were indicated with empty circles.

### Biogeography Analyses

We compiled regional distribution data for each species from peer-reviewed literature and community science platforms, including GBIF, iNaturalist, and Odonata Central. We reconstructed the historical biogeography of Macromiidae using likelihood-based models in BioGeoBEARS (Matzke, 2013). We applied three primary models: DEC (Dispersal-Extinction-Cladogenesis; Ree & Smith, 2008), DIVALIKE (a likelihood implementation of Dispersal-Vicariance Analysis; Ronquist, 1997), and BAYAREALIKE (a Bayesian-like model approximating area inheritance processes; Landis et al., 2013), and further tested each model with the inclusion of the founder-event speciation parameter (+J), which accounts for rare long-distance dispersal or “jump” events (Matzke, 2013). All analyses were performed using our dated phylogeny inferred from our divergence time analyses. Biogeographic regions were defined as in recent odonate biogeographic studies (Abbott et al., 2022; Kalkman et al., 2022; Onsongo et al., in review) and global ecoregion frameworks (Olson et al., 2001), with adjustments to reflect both geographical barriers and phylogenetic distribution of taxa. The regions included the Indo-Malayan region (A - including Sichuan, Hubei, Anhui, and Jiangsu provinces of China), Palearctic (B), Afrotropical realm (C), Wallacea (D - representing the faunal transition zone between Indo-Malaya and Australasia), Nearctic (E - excluding the montane regions of northern and central Mexico), Madagascar and Seychelles (treated separately from mainland Afrotropics due to early isolation and high endemism), and the Australasian realm (F - including Australia, New Guinea, New Zealand, New Caledonia, and the Solomon Islands). Species occurrences were coded according to these regions using a presence/absence matrix. For each model, we set the max range size equal to the largest number of regions occupied by any extant species. Likelihood and ancestral range estimation were performed using the BioGeoBEARS pipeline and marginal probabilities were calculated for each internal node. We compared models using the Akaike Information Criterion (AIC and AICc) calculated from log-likelihoods, to identify the best supported model and scenario of range evolution within the group.

### Ancestral State Reconstruction of habitat preferences and Diversification analyses

We coded habitat preference as a binary trait with lotic environments (rivers and streams) assigned as state 0, and lentic environments (lakes and ponds) as state 1. Taxon names in the dataset were matched to the ultrametric time-calibrated phylogeny of Macromiidae generated using MCMCtree (Yang 2007), and pruned to exclude outgroups using the *ape* package in R. The resulting tree spanned ∼158 MY, providing the temporal framework for all downstream analyses. We estimated ancestral states for habitat preference using marginal reconstructions under the HiSSE framework (Beaulieu & O’Meara 2016). Posterior probabilities of lotic and lentic states were calculated for each internal node, with uncertainty visualized as proportional pies at nodes (Fig 6). Reconstructions were implemented in the R package *hisse* (Beaulieu & O’Meara 2016), with visualization in *ape* (Paradis & Schliep 2019) and *phytools* (Revell 2012). In addition, we assessed whether diversification dynamics were associated with habitat preference using state-dependent speciation and extinction models (SSE). Four model classes were fit (i) CID-2: character-independent null model with two diversification rate classes, (ii) CID-4: character-independent null model with four hidden classes, (iii) BiSSE-like: binary trait-dependent model without hidden states, and (iv) HiSSE: binary trait-dependent model allowing one hidden state (four diversification rate classes). All models were implemented in *hisse* (Beaulieu & O’Meara 2016). Sampling fractions were specified to correct for incomplete sampling coverage of extant taxa: 58/120 species (0.483) for lotic and 4/6 species (0.667) for lentic. Model fit was compared using the corrected Akaike Information Criterion (AICc). Diversification rates were extracted from the best-fitting model and mapped onto branches as continuous gradients.

## RESULTS

### Phylogenetic analyses

Our concatenated phylogenetic analyses recovered a well-resolved tree of Macromiidae with strong overall support across most nodes (Fig. 3). Members of Cavilabiata, including *Cordulegaster bidentata*, *Erythemis guttata*, *Perithemis tenera*, and *Somatochlora alpestris*, are recovered as sister to the rest of the taxa, i.e., Macromiidae. This and other intergeneric outgroup relationships were supported by high bootstrap values (BS=100) and low to intermediate gene and site concordance factors (gCF<40; sCF<70).

**Figure 3:**
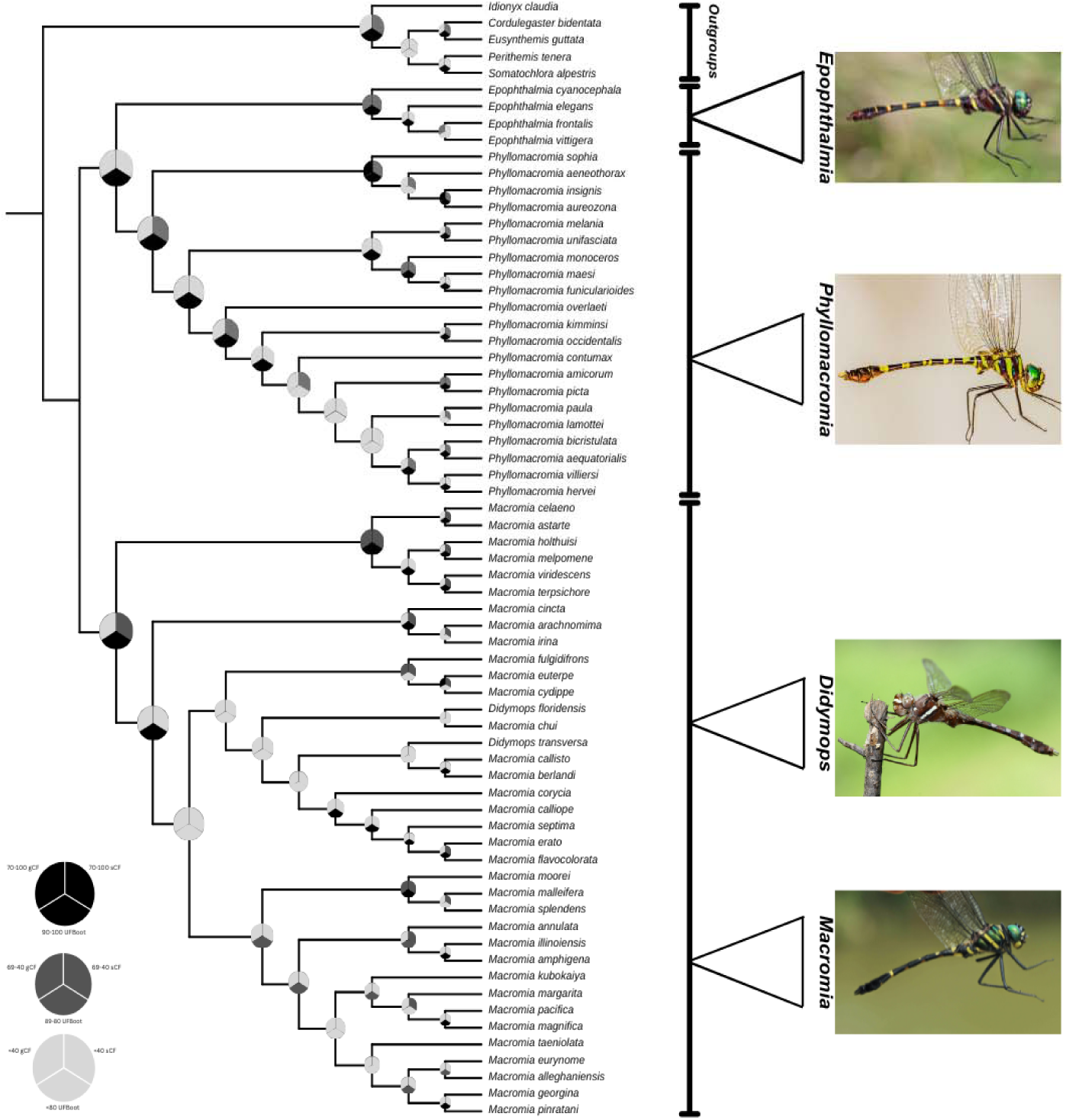
Maximum-likelihood phylogeny of family Macromiidae (Odonata: Anisoptera) from concatenated data in IQ-TREE. Nodal support values represent bootstrap values, gene concordance factors, and the site concordance factors.

Within Macromiidae, three major lineages were recovered: *Epophthalmia*, *Phyllomacromia*, and *Macromia* sensu lato (including *Didymops*). *Epophthalmia* was recovered as sister to *Phyllomacromia*, and this relationship was strongly supported. Within *Epophthalmia*, we recovered *E. cyanocephala* as sister to all of *Epophthalmia* with strong support (BS=100; gCF>70; sCF>70). Both *Epophthalmia* and *Phyllomacromia* were monophyletic and internally well resolved. Within *Phyllomacromia*, multiple well-supported species groups were recovered, including the *P. melania-P. funicularioides* group (BS=100; albeit with low concordance - gCF<40; sCF<40) and the *P. sophia-P. aureozona* group (BS=100; gCF>70; sCF<70), reflecting strong and consistent signal for potential subclades. We recovered *Macromia* sensu lato (including *Didymops*) with strong bootstrap support (BS=100; but with some disconcordance - gCF<40; sCF>40). However, relationships among *Macromia* species suggest that *Didymops,* which is paraphyletic, is nested within the *Macromia*. The two *Didymops* species, *D. floridensis* and *D. transversa*, did not form a monophyletic group; instead, they were recovered in two different parts of the *Macromia* clade. *D. floridensis* recovered as sister to *M. chui*, while *D. transversa* was placed as sister to the *M. callisto + M. berlandi* clade. However, these relationships were not strongly supported; with lower gCF and sCF values relative to BS values, there appears to be gene-tree conflict near the base of *Macromia*. Despite this discordance along the backbone, several species-level relationships within *Macromia* were strongly recovered with respect to bootstrap support. These include the *M. pacifica + M. magnifica* pair (BS=100; with low concordance - gCF<40; sCF<40), the *M. celaeno-M. terpsichore* clade (BS=100; gCF<70; sCF<70), and the *M. illinoiensis* + *M. amphigena* pair (BS=100; with low concordance - gCF<40; sCF<40). Multiple crown clades across *Macromia* were consistently recovered with intermediate bootstrap support, while deeper relationships showed low gene and site concordance, indicating limited agreement among loci, uncertainty in backbone structure of the genus, and the need for broader sampling to improve resolution.

### Morphological analyses, character mapping and ancestral state reconstruction

We assessed the evolutionary history of seven male genitalic characters across Macromiidae and the outgroup taxa using an equal-rates (ER) model, accounting for polymorphism and missing data. Reconstruction estimates varied among characters, influenced by the proportion of observed tip states (coverage) and reflected in our calculated Consistency Index and Retention Index values (Table 2). Nodal support across the different traits were calculated as marginal probabilities of the most likely state, where >0.8 = high, >0.6 <0.8 = moderate, and <0.6 = low. *Epiproct length* had the highest scoring completeness but exhibited substantial homoplasy (Table 2). Three discrete states were found in taxa across the tree, with multiple gains and losses of states inferred across both the ingroups and outgroups (Fig. 4). *Epiproct tip width* was a binary character scored in most taxa (Table 2).

**Figure 4:**
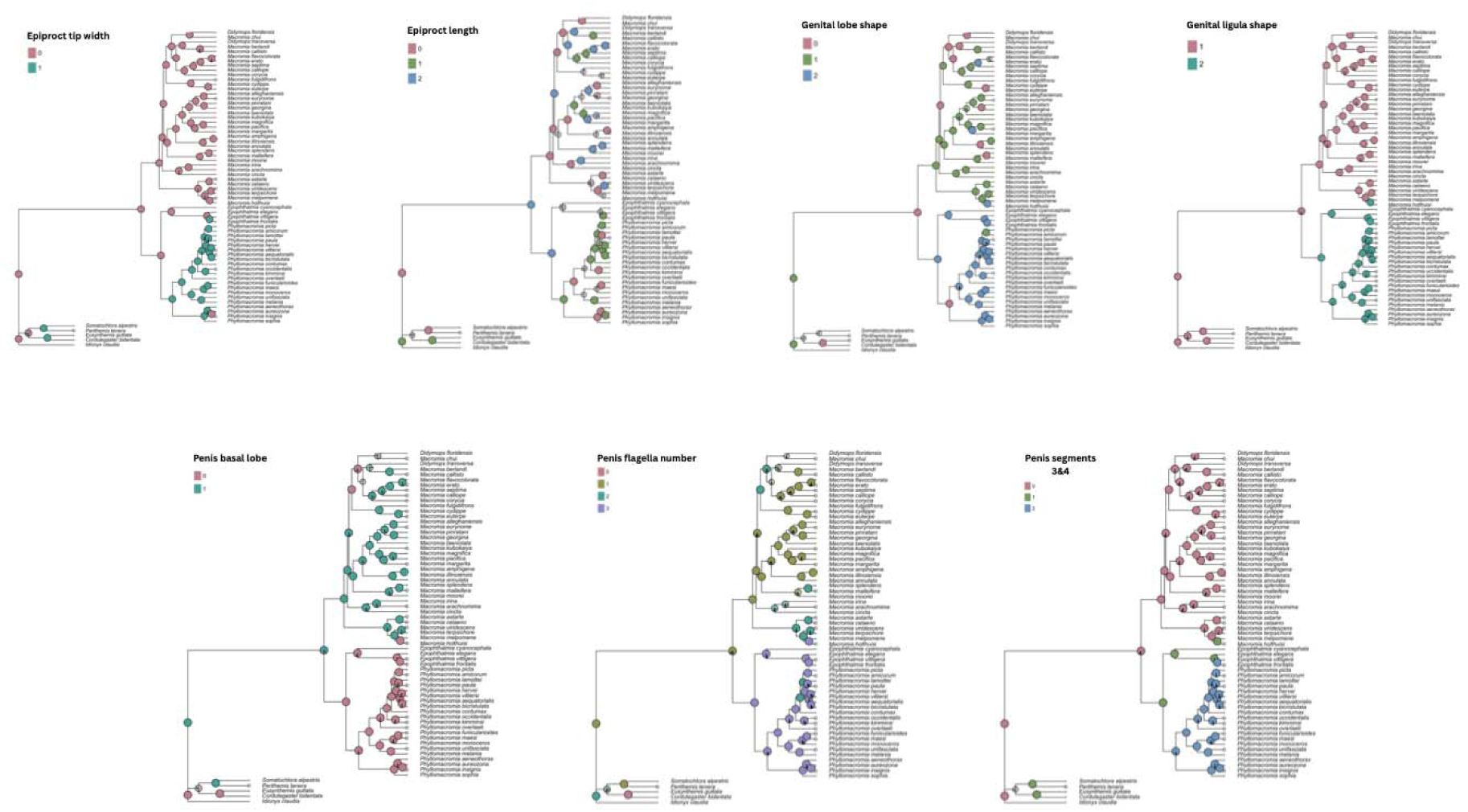
Ancestral state reconstructions of seven male genitalic characters in Macromiidae. Ancestral state estimates were inferred under an ER model using a dated phylogeny. Each panel shows the marginal likelihoods of reconstructed states for a single morphological character: (A) epiproct tip width, (B) epiproct length, (C) genital lobe shape, (D) genital ligula shape, (E) presence of penis basal lobe, (F) number of penis flagella, and (G) shape of penis segments 3&4. Tip states are shown as filled circles and missing data indicated by empty circles. Internal node reconstructions are represented as proportional likelihood pies. Colors correspond to discrete states.

**Table 2:** Genitalic character coding, scoring completeness, and character-fit indices (Consistency Index and Rentention Index) for Ancestral State Reconstruction.

| Character | Number of states | Coverage (%) | CI | RI |
| --- | --- | --- | --- | --- |
| Epiproct length | (3) In relation to cerci - Shorter, Longer, Equal | 93% | 0.09 | 0.34 |
| Epiproct tip width | (2) Narrow, Wide | 85% | 0.33 | 0.89 |
| Genital lobe shape | (3) Pointed, Rounded, Truncate | 90% | 0.14 | 0.63 |
| Genital ligula shape | (3) Flat, Curled, Concave | 67% | 1 | 1 |
| Penis basal lobe | (2) Presence, Absence | 57% | 0.25 | 0.81 |
| Penis flagella number | 1, 2, 3, 4 | 58% | 0.38 | 0.74 |
| Shape of penis segments 3 & 4 | (3) Wide, Intermediate, Slender | 52% | 0.5 | 0.86 |

Reconstructions indicated a predominance of state 0 (narrow) across majority of the family, with state 1 (wide) apparently a unique trait that evolved in a monophyletic subset of *Phyllomacromia;* it showed homoplasy, but relative signal at least in *Phyllomacromia* (Fig. 4)*. Genital lobe shape* was scored as a three-state character and was observed in majority of the taxa. Our reconstructions revealed broad heterogeneity across the family, with all three states distributed across the major lineages. This trait showed substantial homoplasy although retention of phylogenetic signal was moderate (Table 2). *Genital ligula shape* was scored in a smaller number of taxa. Although three states were coded, representing the full range across the ingroups, only two were expressed among the sampled taxa, with curled forms concentrated in *Macromia* + *Didymops* and concave forms found in *Epopthalmia* and *Phyllomacromia* (Fig. 4). *Ligula shape* showed strong phylogenetic structure, with reconstructed states forming distinct clusters corresponding to major clades. Penile characters were generally more difficult to score, but were available for a substantial portion of taxa (Table 2). *Penis basal lobe* presence/absence was observed across ingroup and outgroup taxa, with the basal lobe present in *Macromia* + *Didymops* and absent in *Epophthalmia* and *Phyllomacromia* (Fig. 4). A few internal nodes within *Macromia* show independent losses or reduction of the lobe, with most internal nodes showing moderate to high marginal probabilities. The *Penis flagella number* is also often challenging to score as flagella are prone to breakage, but we were able to score this for over half of the taxa. This was scored as a four-state character, representing the range observed within the superfamily. Our reconstructions revealed a significant state heterogeneity, including multiple events where states 1, 2, or 3 appeared independently across *Macromia* and *Phyllomacromia* (Fig. 4). This is consistent with a structured but moderately homoplastic trait evolution. Finally, the *Shape of penis segments 3&4* was the least consistently scored character (Table 2), but among the taxa we could assess, the wide state was more prevalent within the *Macromia + Didymops* clade, whereas the slender state was concentrated almost exclusively within *Phyllomacromia* (Fig. 4). Our reconstructed node states showed strong concentration of state 0 at deeper nodes with relatively few high-probability transitions.

### Biogeography

Model comparison strongly supported founder event models (Supplementary Table 3). DEC+J and DIVALIKE+J models produced the lowest AICc values (AICc = 153.75) and together accounted for almost all model weight (wAICc = 0.495 each). BAYAREALIKE+J was the next best model (ΔAICc = 7.91), while all non founder event models (DEC, DIVALIKE, BAYAREALIKE) were significantly less supported (ΔAICc > 25). Across all six models, reconstructions at deep nodes differed in the extent of the inferred range. At the root node of Macromiidae, the DEC model recovered the widest ancestral range, with the highest probability state spanning all six regions. Both DEC+J and DIVALIKE+J inferred similarly widespread root states spanning five regions. In contrast, DIVALIKE recovered a much narrower root range with two regions (Indo-Malayan and Afrotropics), and BAYAREALIKE and BAYAREALIKE+J produced the narrowest estimates, with the highest probability states restricted to the Indomalayan region (A). Beyond the root, reconstructions for major clades were largely consistent across models. Nodes within *Phyllomacromia* across all six models were reconstructed with high probability as originating in the Afrotropics. *Epophthalmia* was consistently inferred as having an Indo-Malayan origin, either exclusively or as part of a multi-area origin. Within *Macromia*, internal nodes originated in Indo-Malayan, Australasian, and Nearctic regions across models (Fig. 5). Our examination of node-specific marginal probabilities at Macromiidae crown confirmed the three most widespread root models (DEC, DEC+J, DIVALIKE+J) assigned the highest probability to multi-region ancestral states. In contrast, BAYAREALIKE+J strongly favored a single region state (A, 0.88), and BAYAREALIKE showed low support for all states without a clearly favored state. The DIVALIKE model had relatively moderate support shared among several constricted range states (A, AB, AC) (Supplemental table 4). Despite the differences in the reconstructed ancestral ranges, all six models were consistent in the highest probability regions represented across the major lineages.

**Figure 5:**
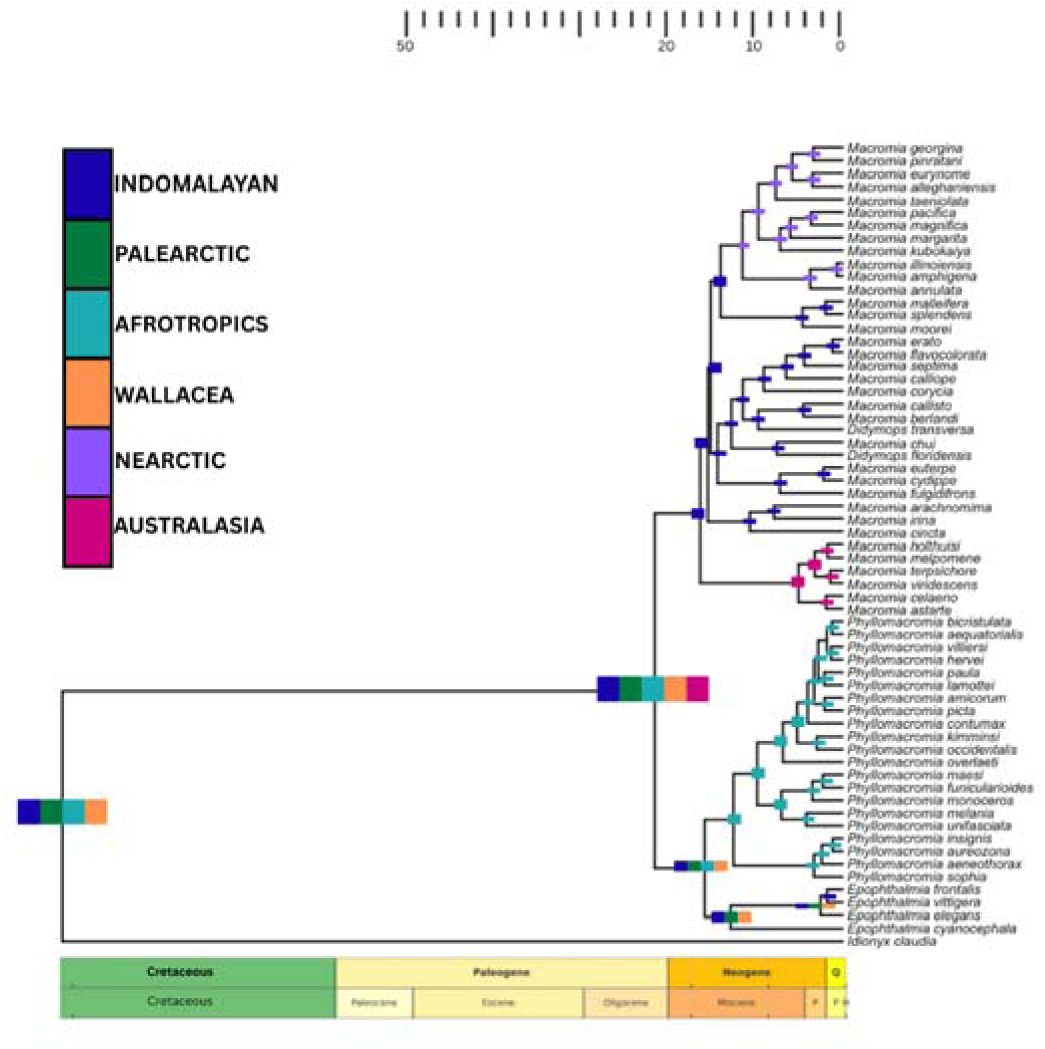
Time-calibrated phylogeny of Macromiidae showing reconstructed ancestral geographic ranges inferred under the best-fitting model (DEC+J). Colors correspond to the seven biogeographic regions used in this study: Indo-Malayan (indigo), Palearctic (green), Afrotropics (teal), Wallacea (orange), Nearctic (purple), and Australasia (mulberry). Ancestral ranges at internal nodes are represented by color blocks indicative of the marginal probability of occupancy for each region.

### Ancestral State Reconstruction of habitat preferences & Diversification

Node-wise marginal reconstructions supported lentic habitat as the ancestral condition for Macromiidae (Fig. 6). The root node showed a high posterior probability for lentic habitat, and this state was also recovered for the stem of the *Macromia* + *Didymops* lineage. In contrast, the common ancestor of *Phyllomacromia* + *Epophthalmia* was reconstructed as being lotic, indicating at least one early transition from lentic to lotic habitat within the family. Among the four models tested (CID-2, CID-4, BiSSE-like, HiSSE), the trait-independent two-class model (CID-2) had the strongest support (AICc = 407.81; Table 3). CID-4 and BiSSE-like were less supported (ΔAICc ∼ 2-3), while the HiSSE model performed the worst (ΔAICc ∼ 22). These results suggest that diversification heterogeneity in Macromiidae is best explained by trait-independent processes rather than trait-dependent models. Branch-specific rate reconstructions showed heterogeneity across the phylogeny (Fig. 6). Diversification rates were summed across hidden state classes within each observed state (lotic vs. lentic), allowing for a direct comparison of diversification ranges between both lineages. Under this framework, lotic and lentic lineages exhibited similar ranges of reconstructed diversification rates (2.98 × 10□¹□ to 0.1446). Warmer-colored branches (indicating higher rates) were concentrated within several derived *Macromia* clades, whereas rates at the backbone showed a wide variation.

**Figure 6:**
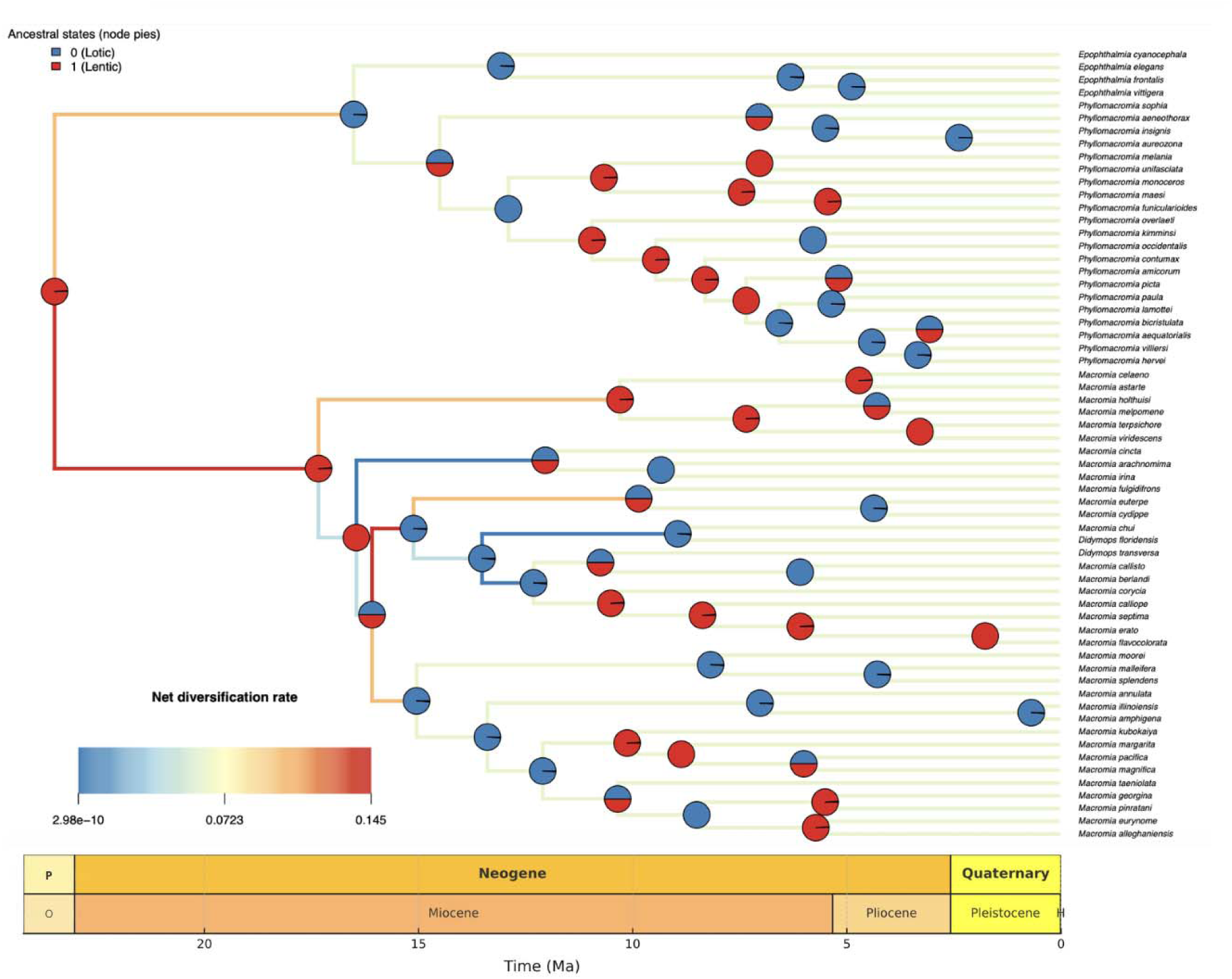
Ancestral state reconstruction of habitat preference across Macromiidae inferred from the best-fit stochastic character mapping model. Node pies represent posterior probabilities of ancestral state for lentic (red) and lotic (blue) habitats, averaged across 200 stochastic maps under an ER model. Branch colors represent net diversification rates, with warmer colors corresponding to higher rates and cooler colors corresponding to lower rates.

**Table 3:** Model selection results for diversification analyses based on HiSSE framework. Four alternative models were fitted: (i) CID-2; (ii) CID-4; (iii) BiSSE-like; and (iv) HiSSE. Models are compared using the sample-size–corrected Akaike Information Criterion (AICc). ΔAICc values are shown relative to the best-fitting model. The CID-2 model received the strongest support (AICc = 407.81), with CID-4 and BiSSE-like models moderately less supported (ΔAICc ∼ 2-3), and the HiSSE model performing much worse (ΔAICc ∼ 22).

| Model | AICc | dAICc |
| --- | --- | --- |
| CID2 | 407.806916 | 0 |
| CID4 | 410.176591 | 2.369675 |
| BiSSE | 411.007896 | 3.200979 |
| HiSSE | 429.637822 | 21.830905 |

Our divergence time estimation recovers a late-Oligocene origin for Macromiidae, estimating the group to be 24 Ma (HPD: 18-33 Ma). The earliest divergence of the *Epophthalmia-Phyllomacromia* lineage from the *Macromia-Didymops* clade, and subsequent splits within each group mostly likely occured within the Miocene (Fig. 7). We estimate the origin of crown *Epopthalmia* to have occurred 14 Ma (HPD: 14-16 Ma), with all internal nodes falling within 0.7-6 Ma, suggesting relatively recent radiation for most intrageneric splits. Within *Epophthalmia*, we recover narrow and young HPDs for sister bifurcations (*E. vittigera* and *E. frontalis*: 0.7-4 Ma). Within *Phyllomacromia*, its divergence is estimated to have occurred 14 Ma (HPD: 8-18 Ma). Several subclades including the *P. picta*, *P. lamottei*, and the *P. funicularioides* groups exhibit internal splits ranging between around 1-5 Ma, which suggests repeated Pliocene diversifications. We estimate the *Macromia* and *Didymops* clade to be 18 Ma (HPD: 16-20 Ma). Early splits involving *Didymops* and the *M. berlandi*-*M. callisto* lineage occurred 14 Ma (HPD: 1-16 Ma).

**Figure 7:**
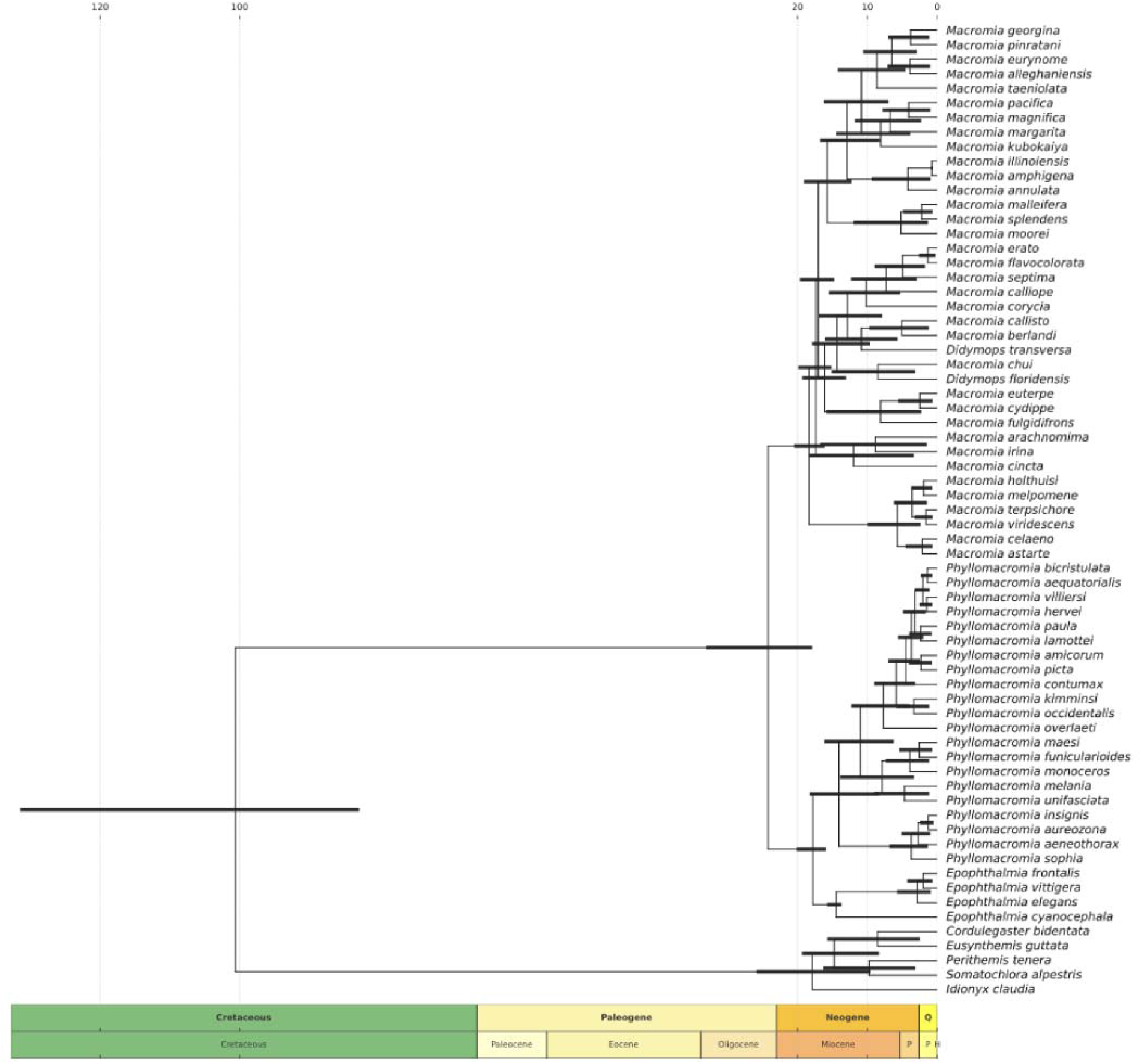
Posterior mean chronogram from MCMCTree (PAML v4.9j) showing estimated divergence times across Macromiidae. Black node bars represent 95% highest posterior density (HPD) intervals for the node ages. Calibrated nodes include the root (>89.5<158.1 Ma) and fossil-based bounds placed on *Epophthalmia*, *Epophthalmia+Phylomacromia*, and several *Macromia* lineages (Table 1). Geological period and epoch boundaries follow the Geologic Time Scale v. 6.0: Geologic Society of America.

## DISCUSSION

### Phylogenetic relationships

Our phylogenetic reconstruction provides new resolution for several relationships within the family and sheds light on the placement of *Didymops* relative to *Macromia*. Previous molecular studies have struggled to recover stable relationships within *Macromia*, in part due to limited taxon sampling, reliance on a small number of loci, or the effects of rapid radiation (Ware et al., 2007; Carle et al., 2015; Bybee et al., 2021; Kosterin et al., 2025). By incorporating broad taxonomic sampling across the group and leveraging high-throughput sampling, our study provides a more resolved framework and reveals areas of phylogenetic discord consistent with earlier hypotheses.

#### Relationship among major genera

Within our ML tree, *Epophthalmia* is recovered as a well-supported, monophyletic lineage. This is in agreement with earlier morphological classifications (Lieftinck, 1932; Lieftinck, 1971) and more modern multilocus analyses, which consistently recognize the genus as distinct from other genera within the family (Carle et al., 2015; Bybee et al., 2021; Kosterin et al., 2025). This distinction was supported by high support values and agreement among loci within the genus. *Phyllomacromia* was also recovered as monophyletic, with robust support across the crown and internal nodes. Previous studies have consistently supported the delimitation of *Phyllomacromia* (May, 1997; Carle et al., 2015; Kosterin et al., 2025), and our results are in agreement with the findings. *Epophthalmia* and *Phyllomacromia* form a strongly supported sister relationship, corresponding to earlier propositions based on morphology (Lieftinck, 1932; 1971; May, 1997). This relationship has been recovered repeatedly in modern molecular approaches, from earlier multilocus studies (Rehn, 2003; Carle et al., 2015) to large-scale targeted enrichment and transcriptomic datasets (Bybee et al., 2021; Kohli et al., 2021; Goodman et al., 2025).

Within *Phyllomacromia*, we recover several subclades that align with historically recognized species groups defined by thoracic patterns, wing venation features, and larval morphology (Lieftinck, 1971; May, 1997; Novelo-Gutierrez & Sites, 2024). Examples include the *P. monoceros-P. maesi* group, the *P. lamottei-P. bicristulata* complex, and East-Central African clusters distinguished by shared diagnostic color patterns and consistent larval traits. Within these groups, we recovered higher concordance among loci than along deeper nodes indicating a stable underlying signal consistent with earlier morphology-based hypotheses.

In contrast to the stability we recovered in the *Phyllomacromia-Epophthalmia* group, the earlier hypothesized relationship between *Macromia* and *Didymops* shows much less phylogenetic stability. We recover crown *Macromia* not forming a clade to the exclusion of *Didymops*. Instead, the two species of *Didymops* are placed in separate positions within *Macromia*; *D. floridensis* recovered as sister to *M. chui*, while *D. transversa* is recovered as sister to the *M. callisto* + *M. berlandi* clade. Although the bootstrap values indicate moderate support for these pairings, the low gene and site concordance show these relationships are not consistently recovered across loci, a common issue with multilocus phylogenetic inference (Jeffroy et al., 2006). The backbone node connecting these *Macromia-Didymops* subclades having a relatively low support further indicates that deeper relationships within the complex remain uncertain. These patterns align with earlier molecular studies involving Macromiidae which also reported limited resolution and relatively low support along the deeper branches of *Macromia* (Ware et al., 2007; Kosterin et al., 2025). Most notably, neither previous work nor the present dataset provides any supported topology in which *Didymops* forms a clade of its own or falls independently outside of *Macromia* indicating that neither represents a monophyletic unit.

Within *Macromia,* several terminal species pairs and subclades in our phylogeny show moderate to high support. This stability is consistent with previous morphological and regional conclusions that have generally reported clear species-level distinctions within the genus (Williamson, 1909; Tenessen, 2019; Kosterin, 2015). However, deeper nodes within the genus show relatively lower support and moderate concordance among loci in our dataset, indicating that the branching order among major lineages of the genus remains uncertain.

Our phylogeny does not recover *Didymops* as monophyletic, and both sampled species appear in separate positions within Macromia. Although the exact placement of each lineage is not heavily supported, no alternative topology in our analyses recovers *Didymops* as a distinct clade or outside *Macromia*. The morphology of the penile characters additionally support *Didymops* and *Macromia* as a single taxon. Given the lack of molecular evidence for the reciprocal monophyly of *Macromia* and *Didymops*, a potential taxonomic approach consistent with our results and conclusions from Kosterin et al. (2025) is to treat *Didymops* as a part of *Macromia*, thereby restoring monophyly through synonymization. This, however, is approached with caution given our limited phylogenetic support.

### Evolution of male genitalic morphology

Comparative genitalic evolution within Macromiidae shows a mixture of strong phylogenetically informative traits and other traits shaped by substantial homoplasy. Although genital structures have been reported to evolve relatively quickly due to sexual selection in Odonata and insects in general (Cordoba-Aguilar, 2005; Eberhard & Leonard, 2010), several traits here retained phylogenetic structure useful for distinguishing major clades within the group.

Across the family, traits associated with the genital ligula, penis structure, and terminal appendages display the strongest phylogenetic structuring. The concave and spoon-shaped genital ligula observed in *Epophthalmia* and *Phyllomacromia* differ from the curved and keeled shape found throughout *Macromia* + *Didymops*, consistent with diagnoses by May (1997). This shift suggests a directional evolution that may reflect changes in mating strategies or female reproductive morphology within the crown clade. Similarly, the transition from wide to more slender terminal structure in penis segments 3 & 4 and the epiproct tip width appears predominant in *Phyllomacromia*, suggesting that the functional complement of abdominal clasping and sperm transfer may have been evolutionarily modified within the lineage. Patterns in penis structure also support a similar grouping, with a well-developed penis basal lobe predominant in *Macromia* + *Didymops* but consistently absent or vestigial in *Epopthalmia* and *Phyllomacromia,* a distinction highlighted by May (1997). Across the different traits, we observe a retention of the ancestral state within the *Macromia* + *Didymops* clade, while *Epophthalmia* and *Phyllomacromia* share derived features that likely evolved after the split between the major lineages. Overall, not all characters showed phylogenetic consistency; a caution of focus is the homoplastic behavior of epiproct length. We observed high variability among closely related taxa, with multiple independent transitions across the tree. This may simply be an artefact of the trait responding more heavily to species-specific mating behavior such as clasping techniques or mate guarding strategies, rather than to evolutionary history. Studies in Odonata and other insects have shown that genitalic components may evolve independently in response to sexual selection, mechanical fit between males and females, and sperm competition (Ebehard & Leonard, 2010; Córdoba-Aguilar, 2003; Cordero-Rivera & Córdoba-Aguilar, 2010). This history of repeated shifts highlights the need to avoid over reliance on any single genitalic trait, especially when those traits are undergoing sexual selection.

### Comparative morphology of the male secondary genitalia support generic relationships

Scanning electron micrographs of the secondary genitalia (Fig. 8) show a pattern of structural similarity within the two major clades within Macromiidae as recognized by May (1997) - *Macromia* + *Didymops* and *Phyllomacromia* + *Epopthalmia.* The overall configuration of the cornua, flagella, and penis segments in *Macromia* and *Didymops* aligns with the conclusion of May (1997) that these two groups only differ in minor details. In both genera, the first penis segment shows sparse setation, a hooked process on segment 2, and comparatively short cornua and flagella in the terminal penis segment, all of which are consistent with diagnosis by May (1997).

**Fig 8:**
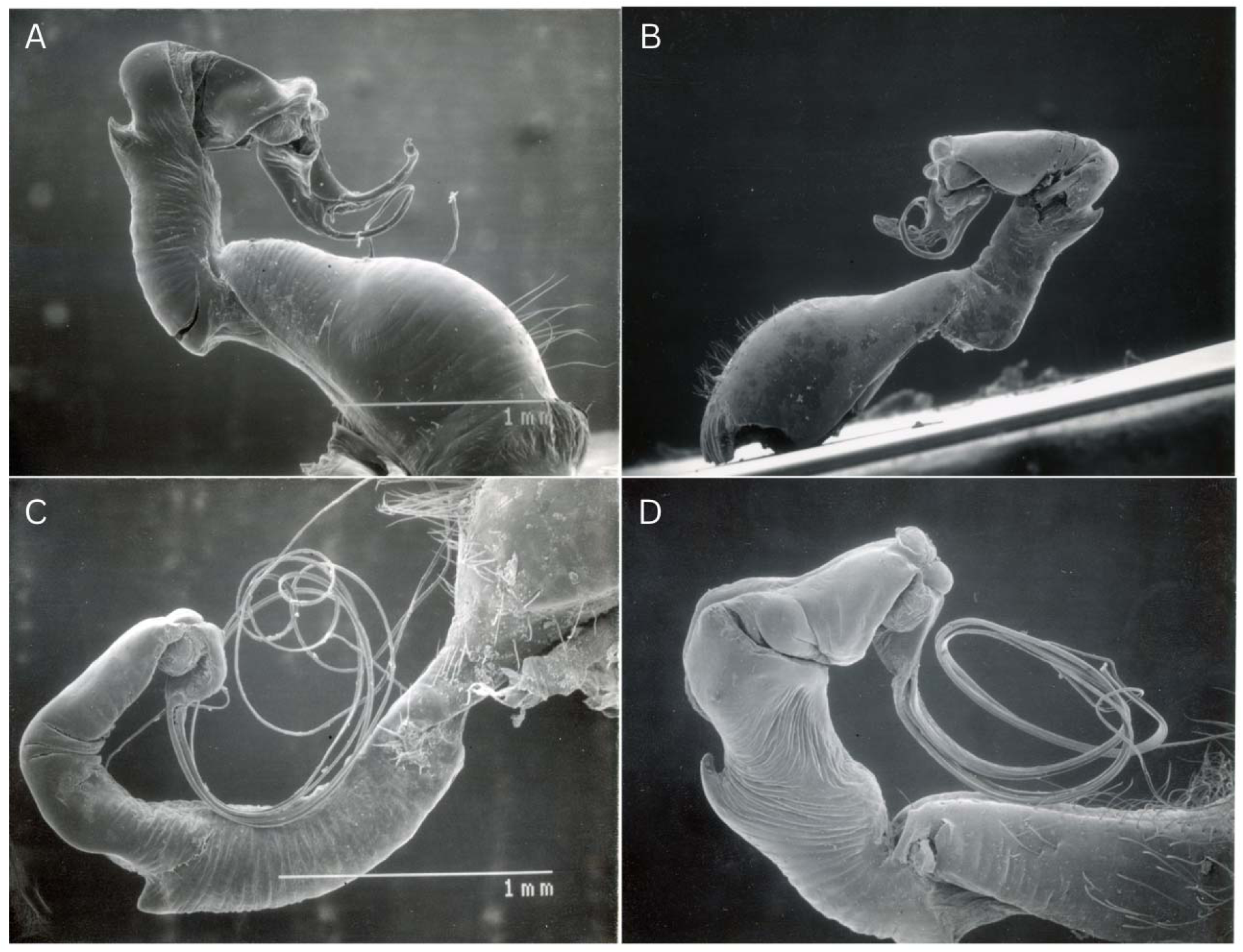
Scanning electron micrographs of the male secondary genitalia from representative individuals of the 4 genera. Showing key differences in setation, penis segment 2 structures, relative length of cornua and flagella. *Didymops* (A) and *Macromia* (B) share sparse setation, a hooked process on segment 2, and short cornua and flagella. *Phyllomacromia* (C) and *Epopthalmia* (D) show dense setation and elongated cornua and flagella.

The SEM images of *Phyllomacromia* and *Epopthalmia* show a different morphological pattern. Both genera exhibit dense setation on the first penis segment, their cornua and flagella are significantly much longer, matching the structures reported in May (1997). Despite these shared features, *Phyllomacromia* shows some distinction from *Epopthalmia* in its penile morphology leading to the hypothesis that *Phyllomacromia* is more different from *Epophthalmia* than *Didymops* is from *Macromia*. Some of these distinctions include the process on penis segment 2 being strongly tooth-like in *Phyllomacromia* but appearing lower and blunter in *Epopthalmia*, both very different from the hooked form seen in *Macromia* and *Didymops.* Similarly, the constriction between penis segments 1 and 2 is pronounced and slender in *Phyllomacromia*, a feature absent in *Epopthalmia*.

### Implications of character indices in lineage diagnosis

Consistency and retention indices indicate that male genitalic characters vary significantly in phylogenetic reliability, with some traits showing pronounced homoplasy while others show signal useful for diagnosing certain lineages. Characters associated with genital ligula shape, penis basal lobe, terminal penile forms, and epiproct tip width exhibit higher retention indices, supporting their interpretation as synapomorphic for the two main lineages in Macromiidae. In contrast, characters associated with epiproct length show both low retention and consistency indices, reflecting repeated gains and losses, and limited informativeness in resolving relationships.

### Biogeography

Our biogeographic analyses identify the DEC+J and DIVALIKE+J as the best fitting models for Macromiidae, consistent with prior work showing that models incorporating founder-event dispersal often outperform purely dispersal-extinction processes (Ree & Smith, 2008; Matzke, 2013). The preference for +J models here does not imply that founder events dominate Macromiidae history, but that including the parameter provides a more accurate probabilities of how ranges changed along the phylogeny. Under our DEC+J model, ancestral Macromiidae is reconstructed as having a widespread old world origin spanning Indomalaya, the Afrotropics, the Palearctic, and Australasia. Although widespread ancestral ranges require cautious interpretation, similar patterns have been inferred in other anisopteran groups with broad modern distributions (Tolman et al., 2024; Goodman et al., 2025), and are consistent with the high dispersal capability and ecological breadth of dragonflies (Bybee et al., 2016). Our reconstructions reveal that most crown diversification occurred during the Paleogene and Neogene, a period in which global climatic and tectonic rearrangements reshaped dispersal pathways (Zhao et al., 2022; Munsterman et al., 2025). Similar temporal patterns have been documented in other odonate lineages (Sanchez-Herrera et al., 2020; Goodman et al., 2025) and similar biogeographic drivers have been suggested (Kalkman et al., 2008; Beatty et al., 2022), suggesting that environmental changes such as Early-Late Miocene Unconformity (EMU-LMU) likely influenced the separation of Macromiidae lineages into different biogeographic regions. The increasingly regional restriction of descendant lineages indicates that once the family dispersed into major continental regions, subsequent range expansions were relatively limited, consistent with niche conservatism (Wang et al., 2025) and the limited intercontinental dispersal seen in dragonflies (Tolman et al., 2025). Our results support the origination of Macromiidae with a wide distribution followed by episodes of range subdivision and regional isolation throughout the Cenozoic. The best fitting models indicate that long distance dispersal events likely contributed to the geographic shaping of extant lineages, while the overall pattern of region-specific colonizations suggests long-term ecological stability influence and deep time barriers in shaping present diversity of the family. Further expansion of geographic sampling, particularly from undersampled regions such as Wallacea and the Afrotropics, will better refine these inferences and clarify the extent to which the different events influence geographical isolation within the family.

### Diversification with respect to habitat preference

Diversification dynamics within Macromiidae show clear heterogeneity across the tree, with several clades exhibiting higher net diversification rates during the Miocene (Fig. 6). This pattern is broadly consistent with diversification trends documented across Odonata (Letsch et al., 2016; Padilla-Morales et al., 2021) and trends within freshwater insects more generally during the Neogene, when tectonic and climatic shifts reorganized global freshwater systems, providing opportunities for merolimnic taxa to colonize new niches (Luo et al., 2020). Previous large-scale Odonata studies suggested that lentic lineages may diversify more rapidly than lotic lineages, with transitions into lentic habitats linked to increased speciation (Letsch et al., 2016; Padilla-Morales et al., 2021). Our ancestral state reconstructions recover lentic habitat use as the ancestral preference, accompanied by relatively high early diversification rates before an eventual cool with subsequent transitions. Within the family, however, our results reveal that lentic vs lotic habitat preference does not directly influence diversification rates. The trait-independent CID2 model was strongly favored over trait-dependent models (BiSSE and HiSSE) and the more complex trait-independent model (CID4). This indicates that the diversification heterogeneity we recover is more accurately explained by background processes independent of the ecological traits in focus (lentic/lotic). Our findings align with critiques of trait-dependent diversification models, which caution against over-interpreting associations between ecological traits and diversification, particularly when background rate heterogeneity may provide a better explanation (Rabosky & Goldberg, 2015; Beaulieu & O’Meara, 2016). We believe that while shifts between lentic and lotic preferences have occurred repeatedly within Macromiidae, these transitions do not necessarily correspond to major changes in speciation or extinction dynamics. While we make these conclusions, we acknowledge some points of consideration; first, binary habitat coding may be an oversimplification of ecological dynamics, as many species utilize different flow gradients or may shift habitats through development (Nakazawa, 2015). Second, SSE models are sensitive to incomplete sampling and extinction estimates, which may obscure subtle trait effects (Rabosky & Goldberg, 2015). Lastly, unobserved traits not captured by the CID2 model, such as dispersal ability, life-history strategies, or range size, may be more accurate drivers of diversification heterogeneity than habitat category alone.

### Divergence time estimation

Our divergence estimations indicate that crown Macromiidae originated in the late Oligocene (∼24 Ma), with subsequent splitting concentrated in the Miocene-Pliocene. This pattern is consistent with genomic and transcriptomic odonate studies, which recover deep Jurassic-Cretaceous origins for major anisopteran lineages but younger Cenozoic crown radiations within derived families (Kohli et al., 2021; Huang et al., 2025). Compared with early diverging families such as Petaluridae, whose extant lineages date back to the Jurassic (Tolman et al., 2024), Macromiidae more closely aligns with Miocene radiations such as *Somatochlora* (Corduliidae), which diversified around 19 Ma (Goodman et al., 2025). The Miocene crown ages estimated for *Epophthalmia* and *Phyllomacromia* (∼14 Ma) and the *Macromia* + *Didymops* clade (∼18 Ma) follow a well-documented Miocene diversification burst driven by climatic oscillations, hydrologic and topographic change (Steinthorsdottir et al., 2020; Dagallier et al., 2024), which odonates are well-suited to capitalize on. Recent work has shown that fossil placement can strongly influence odonate node ages (Kohli et al., 2021; Huang et al., 2025), and as such, absolute ages should be interpreted cautiously. However, the estimated timing of a young Miocene Macromiidae crown recovered here accompanied by rapid Pliocene radiations, is congruent with emerging phylogenomic studies for Odonates.

## Conclusion

Our study provides a phylogenetic framework for Macromiidae, integrating subgenomic-scale data, fossil-calibrated divergence estimates, biogeographic inference, and a focused assessment of male genitalic morphology. Our results strongly support the sister relationships of *Epophthalmia* + *Phyllomacromia* and *Macromia* + *Didymops*, with major lineages diversifying during the Paleogene-Neogene. Although the groups span lentic and lotic habitats, these ecological preferences do not appear to drive diversification directly. Ancestral range estimates reveal an Old-World origin, followed by independent dispersal events shaping present-day distributions. We recover certain genitalic traits reflecting deeper evolutionary history, with *Macromia* + *Didymops* retaining more ancestral forms and *Epophthalmia* + *Phyllomacromia* sharing derived features. The SEM images support this pattern, showing close structural similarity between *Didymops* and *Macromia* and between *Epophthalmia* and *Phyllomacromia*, particularly in setation, segment 2 process, and the relative length of the cornua and flagella. Our analyses also reveal multiple independent state shifts in certain characters, limiting phylogenetic signal and highlighting the need for integrating molecular data alongside morphology. Overall, we present a picture of Macromiidae evolution, providing a foundation for future taxonomic refinement, hypothesis testing, and comparative study on reproductive morphology within the group.

