## Supplemental Table 1 for "Systematics, diversification, and biogeography of Macromiidae (Odonata: Anisoptera)"

**Supplementary Table 1**: Specimen information, locality, date collected, and collector

| Species | Sequencing code | Country | State/Province | Locality | Date | Collector |
| --- | --- | --- | --- | --- | --- | --- |
| *Didymops floridensis* | GEODE17616 | USA | Alabama | Covington Co.; Conecuh NF | 3/28/02 | T. W. & A. J. Donnelly |
| *Didymops transversa* | GEODE17617 | USA | Alabama | Blount County, Mill Creek, near Blount Springs | 5/10/92 | K. J. Tennessen |
| *Epophthalmia elegans* | OD1101 | China |  |  |  |  |
| *Epophthalmia frontalis* | RMNH4010129291 | Cambodia |  |  | 2014 |  |
| *Epophthalmia cyanocephala* | GEODE7611 | India | South Malabar | Walayar Forest | 9/8/47 | P. S. Nathan |
| *Epophthalmia vittigera* | RMNH30105550 | Malaysia | Borneo | Sabah, Ranau-Lohan Road | 1/9/99 | T. W. & A. J. Donnelly |
| *Macromia alleghaniensis* | GEODE17619 | USA | Alabama | Lauderdale County | 7/22/78 | K. J. Tennessen |
| *Macromia amphigena* | GEODE17620 | Japan | Hokkaido Prefecture | Nishioka, Sapporo-city | 6/24/95 | T. K. Yokoyama |
| *Macromia annulata* | GEODE17621 | USA | Texas | San Saba County | 8/2/95 | T. Donnelly |
| *Macromia arachnomima* | RMNH25919144 | Malaysia |  |  | 2008 |  |
| *Macromia astarte* | GEODE17622 | Papua New Guinea | Morobe | Gurakor, Wau Road | 10/25/72 | T. Donnelly |
| *Macromia berlandi* | GEODE17623 | Hong Kong | Sha Lo Tung |  | Jun-98 |  |
| *Macromia calliope* | RMNH4000479250 | China |  |  | 2009 |  |
| *Macromia callisto* | GEODE17624 | Malaysia | Perak Sungai | Ayer Jada | 5/12/98 | A. Donnelly |
| *Macromia celaeno* | GEODE2761 |  |  |  |  |  |
| *Macromia chui* | GEODE17625 | China | Guangdong | Conghua city | 5/15/10 | Zhang Haomiao |
| *Macromia cincta* | RMNH4002185222 | Malaysia |  |  | 2012 |  |
| *Macromia cydippe* | RMNH4002184840 | Malaysia |  |  | 2012 |  |
| *Macromia erato* | GEODE2762 |  |  |  |  |  |
| *Macromia eurynome* | GEODE17630 | Papua New Guinea | Morobe | Bulolo | Aug-84 | S. W. Dunkle |
| *Macromia euterpe* | RMNH4002185034 | Malaysia |  |  | 2010 |  |
| *Macromia flavocolorata* | GEODE17631 | Thailand | Tak Province | Um Phang District, Ya Mo Ki Village | 5/4/98 | T. Donnelly |
| *Macromia fulgidifrons* | RMNH4000479269 | China |  |  | 2011 |  |
| *Macromia holthuisi* | RMNH4003832097 | Indonesia |  |  | 2006 |  |
| *Macromia illinoiensis* | OD1087 |  |  |  |  |  |
| *Macromia irina* | GEODE2764 |  |  |  |  |  |
| *Macromia kubokaiya* | GEODE17632 | Japan | Ryukyus, Okinawa | Kunigami, Yona, Okuni-rindo | 7/6/08 | Y. Kishida |
| *Macromia magnifica* | GEODE17633 | Canada | British Columbia | Cultus Lake | 7/14/71 | D. R. & M. L. Paulson |
| *Macromia malleifera* | RMNH4000479280 | China |  |  | 2011 |  |
| *Macromia margarita* | GEODE17634 | USA | North Carolina | Macon County, Cullasaja River | 7/8/82 | K. J. Tennessen |
| *Macromia melpomene* | RMNH4003831975 | Indonesia |  |  | 2006 |  |
| *Macromia moorei* | RMNH4000479238 | China |  |  | 2008 |  |
| *Macromia pacifica* | GEODE17635 | USA | Missouri | Gasconade County, Bourbeuse Road | 6/18/02 | T. W. & A. J. Donnelly |
| *Macromia pinratani* | GEODE17636 | Thailand | Ranong | Muang Shone | Mar-96 | Pinratana |
| *Macromia septima* | RMNH4003831547 | Cambodia |  |  | 2011 |  |
| *Macromia splendens* | RMNH.INS.507575 | France | Mialet | 44.106, 3.948 | 6/18/12 | Keijl, T. O. |
| *Macromia taeniolata* | GEODE17637 | USA | Alabama | Lauderdale County, Tennessee River | 9/7/79 | K. J. Tennessen |
| *Macromia terpsichore* | RMNH4003831969 | Indonesia |  |  | 2006 |  |
| *Macromia viridescens* | GEODE2765 |  |  |  |  |  |
| *Phyllomacromia aeneothorax* | RMNH30105869 | Liberia |  |  | 2011 |  |
| *Phyllomacromia aequatorialis* | RMNH4010122260 | Angola |  |  | 2013 |  |
| *Phyllomacromia amicorum* | RMNH30232991 | Liberia |  |  | 2010 |  |
| *Phyllomacromia aureozona* | RMNH30231575 | Democratic Republic of Congo |  |  | 2010 |  |
| *Phyllomacromia bicristulata* | RMNH4012544345 | Gabon |  |  | 2013 |  |
| *Phyllomacromia contumax* | RMNH30231688 | Democratic Republic of Congo |  |  | 2010 |  |
| *Phyllomacromia funicularioides* | RMNH30232992 | Liberia |  |  | 2010 |  |
| *Phyllomacromia hervei* | RMNH4000463203 | Angola |  |  | 2012 |  |
| *Phyllomacromia insignis* | RMNH4010122704 | Gabon |  |  | 2013 |  |
| *Phyllomacromia kimminsi* | RMNH4003826962 | Zambia |  |  | 2010 |  |
| *Phyllomacromia lamottei* | RMNH30108065 | Liberia |  |  | 2012 |  |
| *Phyllomacromia maesi* | RMNH4010122770 | Gabon |  |  | 2013 |  |
| *Phyllomacromia melania* | RMNH4003826927 | Zambia |  |  | 2010 |  |
| *Phyllomacromia monoceros* | RMNH4002191053 | Democratic Republic of Congo |  |  | 2011 |  |
| *Phyllomacromia occidentalis* | RMNH30108089 | Liberia |  |  | 2012 |  |
| *Phyllomacromia overlaeti* | RMNH4003831771 | Cameroon |  |  | 2008 |  |
| *Phyllomacromia paula* | RMNH30230406 | Democratic Republic of Congo |  |  | 2010 |  |
| *Phyllomacromia picta* | GEODE7715 | Rwanda | N. U. of Rwanda University Arboretum | S2.612869, E29.75995 | 5/7/13 | Bybee, S.M., G. Svenson, N. Hardy |
| *Phyllomacromia sophia* | RMNH30232954 | Liberia |  |  | 2010 |  |
| *Phyllomacromia unifasciata* | RMNH4003826919 | Zambia |  |  | 2010 |  |
| *Phyllomacromia villiersi* | RMNH4012544536 | Gabon |  |  | 2014 |  |
| *Cordulegaster bidentata* | RMNH4000482741 | France |  |  | 2012 |  |
| *Eusynthemis guttata* | GEODE17891 |  |  |  |  |  |
| *Perithemis tenera* | GEODE7739 |  |  |  |  |  |
| *Somatochlora alpestris* | GEODE17828 | Norway | Soitun | Troms | 7/25/04 | Hjalmar Westgard |
| *Idionyx claudia* | GEODE17864 |  |  |  |  |  |

**Supplementary Table 2**: Morphology matrix including all ingroups and outgroups used in this study

| Species | Genital ligula | Penis segments 3 & 4 | Penis flagella number | Penis basal lobe | Gen lobe shape | Epiproct length | Epiproct tip width |
| --- | --- | --- | --- | --- | --- | --- | --- |
| *Didymops floridensis* Davis, 1921 | 1 | 0 | 1 | 1 | 1 | 0 | 0 |
| *Didymops transversa* (Say, 1840) | 1 | 0 | 1 | 1 | 1 | 0 | 0 |
| *Epophthalmia elegans* (Brauer, 1865) | 2 | 1 | 3 | 0 | 2 | 2 | 0 |
| *Epophthalmia frontalis* Selys, 1871 | ? | ? | 3 | ? | 2 | 0,2 | 0 |
| *Epophthalmia cyanocephala* Burmeister, 1839 | 2 | 1 | 3 | 0 | 2 | 1,2 | 0 |
| *Epophthalmia vittigera* (Rambur, 1842) | 2 | 1 | ? | ? | 2 | 0,1 | 0 |
| *Macromia alleghaniensis* Williamson, 1909 | 1 | 0 | 1 | 1 | 2 | 2 | 0 |
| *Macromia amphigena* Selys, 1871 | 1 | 0 | 1 | 1 | 1 | 0 | 0 |
| *Macromia annulata* Hagen, 1861 | 1 | 0 | 1 | 1 | 1 | 1 | 0 |
| *Macromia arachnomima* Lieftinck, 1953 | ? | ? | ? | ? | 1 | 2 | 0 |
| *Macromia astarte* Lieftinck, 1971 | 1 | ? | ? | ? | 1 | 0 | ? |
| *Macromia berlandi* Lieftinck, 1941 | 1 | 0 | 1 | 1 | 1 | 0 | 0 |
| *Macromia calliope* Ris, 1916 | 1 | ? | ? | ? | 0 | 2 | 0 |
| *Macromia callisto* Laidlaw, 1922 | 1 | ? | ? | ? | 0 | 2 | 0 |
| *Macromia celaeno* Lieftinck, 1955 | 1 | ? | ? | ? | 1 | 2 | 0 |
| *Macromia chui* Asahina, 1968 | ? | ? | ? | ? | 0 | 2 | 0 |
| *Macromia cincta* Rambur, 1842 | 1 | 0 | 2 | 1 | 0 | 2 | 0 |
| *Macromia cydippe* Laidlaw, 1922 | ? | ? | ? | ? | 1 | 1 | 0 |
| *Macromia erato* Lieftinck, 1950 | 1 | ? | ? | ? | 0 | 0 | 0 |
| *Macromia eurynome* Lieftinck, 1942 | ? | ? | ? | ? | ? | 2 | ? |
| *Macromia euterpe* Laidlaw, 1915 | ? | ? | ? | ? | 1 | 1 | 0 |
| *Macromia flavocolorata* Fraser, 1922 | 1 | 0 | 2 | 0 | 1 | 0 | 0 |
| *Macromia fulgidifrons* Wilson, 1998 | 1 | 0 | 1 | 1 | 1 | 1 | ? |
| *Macromia holthuisi* Kalkman, 2008 | 1 | ? | ? | ? | 1 | 2 | 0 |
| *Macromia illinoiensis* Walsh, 1862 | 1 | ? | 1 | 1 | 0 | ? | 0 |
| *Macromia irina* Lieftinck, 1950 | 1 | 0 | 1 | 1 | 1 | 0 | 0 |
| *Macromia kubokaiya* Asahina, 1964 | 1 | ? | ? | ? | 1 | 0 | 0 |
| *Macromia magnifica* Selys, 1874 | 1 | 0 | 1 | 1 | 1,2 | 1 | 0 |
| *Macromia malleifera* Lieftinck, 1955 | 1 | ? | ? | ? | 1 | 0,2 | 0 |
| *Macromia margarita* Westfall, 1947 | ? | ? | ? | ? | 1 | 2 | 0 |
| *Macromia melpomene* Ris, 1913 | 1 | 0 | 2 | 1 | 1 | 0 | ? |
| *Macromia moorei* Selys, 1874 | 1 | 0 | 1 | 1 | 0 | 1,2 | 0 |
| *Macromia pacifica* Hagen, 1861 | 1 | 0 | 1 | 1 | ? | 0,2 | 0 |
| *Macromia pinratani* Asahina, 1983 | 1 | 0 | 1 | 1 | 1 | 0,2 | 0 |
| *Macromia septima* Martin, 1904 | 1 | ? | ? | ? | 0 | 0 | 0 |
| *Macromia splendens* (Pictet, 1843) | 1 | 0 | 1 | 1 | 2 | 2 | 0 |
| *Macromia taeniolata* Rambur, 1842 | ? | 0 | 1 | 1 | 1 | 0,1 | 0 |
| *Macromia terpsichore* Förster, 1900 | 1 | 0 | 2 | 1 | 1 | 0 | 0 |
| *Macromia viridescens* Tillyard, 1911 | 1 | ? | ? | ? | 1 | ? | ? |
| *Phyllomacromia aeneothorax* (Nunney, 1895) | 2 | 2 | 3 | 0 | 2 | 1 | 1 |
| *Phyllomacromia aequatorialis* Martin, 1906 | ? | ? | 2 | 0 | 2 | 1 | ? |
| *Phyllomacromia amicorum* (Gambles, 1979) | ? | ? | ? | ? | 0 | 1 | 1 |
| *Phyllomacromia aureozona* (Pinhey, 1966) | ? | 2 | 3 | 0 | 2 | 0 | 1 |
| *Phyllomacromia bicristulata* (Legrand, 1975) | 2 | ? | 3 | 0 | 1 | 1 | 1 |
| *Phyllomacromia contumax* Selys, 1879 | 2 | 2 | 3 | 0 | 2 | 0,1 | 1 |
| *Phyllomacromia funicularioides* (Legrand, 1983) | ? | ? | ? | ? | 2 | 1 | 1 |
| *Phyllomacromia hervei* (Legrand, 1980) | ? | 2 | 3 | 0 | 2 | 1 | 1 |
| *Phyllomacromia insignis* (Kirby, 1889) | ? | ? | ? | ? | 2 | 0 | ? |
| *Phyllomacromia kimminsi* (Fraser, 1954) | ? | ? | ? | ? | 2 | 1 | 1 |
| *Phyllomacromia lamottei* (Legrand, 1993) | ? | 2 | 3 | 0 | ? | 0 | 1 |
| *Phyllomacromia maesi* (Schouteden, 1917) | ? | ? | ? | ? | 2 | 0 | 1 |
| *Phyllomacromia melania* (Selys, 1871) | 2 | 2 | 3 | 0 | 2 | 0 | 1 |
| *Phyllomacromia monoceros* (Förster, 1906) | 2 | 2 | 3 | 0 | 2 | 0,1 | 1 |
| *Phyllomacromia occidentalis* (Fraser, 1954) | ? | ? | ? | ? | ? | 0 | ? |
| *Phyllomacromia overlaeti* (Schouteden, 1934) | ? | ? | ? | ? | 2 | 1 | 1 |
| *Phyllomacromia paula* (Karsch, 1892) | ? | ? | ? | ? | 2 | ? | 1 |
| *Phyllomacromia picta* (Hagen in Selys, 1871) | 2 | 2 | 3 | 0 | 2 | 1 | 1 |
| *Phyllomacromia sophia* (Selys, 1871) | 2 | 2 | 3 | 0 | 2 | 1 | 0 |
| *Phyllomacromia unifasciata* (Fraser, 1954) | ? | ? | ? | ? | 2 | 0,1 | 1 |
| *Phyllomacromia villiersi* (Legrand, 1992) | 2 | 2 | 2 | 0 | ? | 1 | ? |
| *Cordulegaster bidentata* | 1 |  | 0 | 0 | 0 | 1 | 1 |
| *Eusynthemis guttata* | 1 |  | 1 | 1 | ? | 0 | 1 |
| *Perithemis tenera* | ? | ? | ? | ? | ? | ? | ? |
| *Somatochlora alpestris* | 1 |  | 2 | 0 | 1 | 1 | 0 |
| *Idionyx claudia* | 1 |  | 2 | 1 | 1 | 1 | 0 |

**Supplementary Table 3**: BioGeoBears model comparison information showing AIC, AICc, and ΔAICc values

| Model | LogLik | k | AIC | AICc | Delta_AICc | AICc_weight |
| --- | --- | --- | --- | --- | --- | --- |
| DECJ | -73.672 | 3 | 153.343 | 153.75 | 0 | 0.495 |
| DIVALIKEJ | -73.672 | 3 | 153.343 | 153.75 | 0 | 0.495 |
| BAYAREALIKEJ | -77.628 | 3 | 161.255 | 161.662 | 7.912 | 0.009 |
| DEC | -88.175 | 2 | 180.35 | 180.55 | 26.8 | 0 |
| DIVALIKE | -99.342 | 3 | 204.683 | 205.09 | 51.34 | 0 |
| BAYAREALIKE | -106.622 | 2 | 217.245 | 217.445 | 63.695 | 0 |

**Supplementary Table 4**: Node-wise marginal probabilities of the six biogeographic models tested; Node 65 (root node)

| **State** | **DEC** | **DEC_J** | **DIV** | **DIV_J** | **BAY** | **BAY_J** |
| --- | --- | --- | --- | --- | --- | --- |
| **A** | 0.171085886621419 | 0.114099383803798 | 0.0281730193382139 | 0.114101999631688 | 0.0114436597834197 | 0.876695015348144 |
| **AF** | 0.0638940072385067 | 0.0382786797281472 | 0.0229161959867259 | 0.0382874523988877 | 0.0165204437899911 | 0.0205022754568556 |
| **AC** | 0.0596243188116051 | 0.0606503857361118 | 0.02525448478952 | 0.0606623153639558 | 0.0180516525438794 | 0.0204911913665273 |
| **AB** | 0.0435932004083604 | 0.0317923094253146 | 0.0226707275906025 | 0.0317908897636005 | 0.0162995407500252 | 0.0198704049685098 |
| **AD** | 0.0435834230953326 | 0.0424343424003715 | 0.0229182393695866 | 0.0424305959311283 | 0.0165925534906587 | 0.0283721984044831 |
| **ACF** | 0.0343876788360917 | 0.0337025883006302 | 0.0223755230414034 | 0.0337086761301659 | 0.0217500584204529 | 0.000541748413860095 |
| **AE** | 0.0341000871251686 | 0.00075619482002919 | 0.0218187266537315 | 0.000756639654193133 | 0.0169554412522812 | 0.0197952742275835 |
| **ABF** | 0.0298679016490531 | 0.018199649923672 | 0.0206733757451595 | 0.018200072617956 | 0.0200494653502265 | 0.000548969236085871 |
| **ADF** | 0.0292130369125968 | 0.034728472068504 | 0.0208468677877272 | 0.0347294792480037 | 0.0203363464820864 | 0.000757113482043058 |
| **ABC** | 0.0285635092418332 | 0.0731390534668831 | 0.0222973536041448 | 0.0731414680047428 | 0.0215239227807795 | 0.000547070760224788 |
| **ACD** | 0.0282517572443557 | 0.081168250634716 | 0.0224723185963499 | 0.0811696599027074 | 0.0218194600856478 | 0.00076411165807473 |
| **AEF** | 0.0260578067223484 | 0.00148641362092838 | 0.0200748640847104 | 0.00148761211805297 | 0.020691822005694 | 0.000547067282299066 |
| **ABD** | 0.02374367855456 | 0.0574297256522201 | 0.0207356426825144 | 0.0574233237587994 | 0.02011877344509 | 0.000786180865217855 |
| **ABCF** | 0.0226852017127194 | 0.0494253735353054 | 0.0206333530254202 | 0.0494269720139381 | 0.0241442424893558 | 1.74838881341572E-05 |
| **ACDF** | 0.02196963433103 | 0.0598390132432318 | 0.0207675949772857 | 0.0598408401720816 | 0.0244235847880467 | 2.28731329895926E-05 |
| **ACE** | 0.0206929217241857 | 0.000381457678037087 | 0.0216312951514132 | 0.000381674814058867 | 0.0221932827330545 | 0.000545089167833738 |
| **ABDF** | 0.0200899289615344 | 0.041026457445279 | 0.019498545758162 | 0.0410235295629224 | 0.0228235486339384 | 2.77638131401064E-05 |
| **ABCD** | 0.0200633253431879 | 0.153374936134931 | 0.0207432685445009 | 0.153360874289464 | 0.0242090078363666 | 2.79820695582593E-05 |
| **ABE** | 0.0196923056098495 | 0.000143406387207565 | 0.0199428074544889 | 0.000143521947119519 | 0.0204732050173535 | 0.000550911392336107 |
| **ABEF** | 0.0187060864195476 | 0.000289612800328532 | 0.0188888932422734 | 0.000289995175098911 | 0.0231596532642389 | 2.18407564236238E-05 |
| **ACEF** | 0.0185024687937731 | 0.000864235157791646 | 0.0201254411581245 | 0.000864931513466261 | 0.0247763320943453 | 1.74362619203768E-05 |
| **ADE** | 0.017460543232955 | 0.000241762402439545 | 0.0201038785144776 | 0.000241835696664529 | 0.0207629372922517 | 0.000773569729706073 |
| **ABCDF** | 0.0169446335257587 | 0.0982201513229533 | 0.0195839607685917 | 0.0982126297404046 | 0.0260609389919623 | 8.86957753970451E-07 |
| **ADEF** | 0.0164758277080995 | 0.000550246124676026 | 0.0190135154939613 | 0.000550524158384755 | 0.023434003310544 | 2.7393660895014E-05 |
| **ABCE** | 0.0155201142924646 | 0.000170945735629335 | 0.0200843405070596 | 0.00017098638631693 | 0.0245608157699268 | 2.17538524914931E-05 |
| **ABCEF** | 0.0148399024798273 | 0.000166257361815172 | 0.0190544072539867 | 0.000166504170529783 | 0.0263886791685228 | 8.41244410850369E-07 |
| **ABDE** | 0.0140501482906655 | 0.000143587813605 | 0.0189504799447047 | 0.000143607850541872 | 0.0232258845464413 | 3.24979590013036E-05 |
| **ACDE** | 0.0139928346223594 | 0.000242054814051539 | 0.0202101491527282 | 0.000242065108114123 | 0.0248426679551534 | 2.76009861631486E-05 |
| **ABDEF** | 0.0137302761011921 | 0.000143554895642096 | 0.0181769202712527 | 0.000143744191312361 | 0.0251505235189404 | 1.86626486687494E-06 |
| **ACDEF** | 0.0133168278913544 | 0.000329352200311683 | 0.0191566192464636 | 0.000329531569966007 | 0.0266513879647774 | 8.75778205357559E-07 |
| **ABCDE** | 0.0120595642852925 | 0.000244477667930248 | 0.0191475900651861 | 0.000244395177360875 | 0.0264497824296233 | 1.87975725681685E-06 |
| **ABCDEF** | 0.0116047149542075 | 8.66127503825553E-05 | 0.018367813205219 | 8.67436943569332E-05 | 0.0277448747084915 | 2.14532374900927E-08 |
| **F** | 0.00336117281761509 | 0.000450233090565545 | 0.00679244030076871 | 0.000449840904872475 | 0.00253903094187204 | 0.001618943770703 |
| **C** | 0.00312252770883767 | 0.00065605218717317 | 0.00834806057564838 | 0.000655521093491836 | 0.00315165012040786 | 0.00160716456681351 |
| **CF** | 0.00280940108811088 | 0.000259297740566049 | 0.0111831016436669 | 0.00025905611202329 | 0.00773075967904489 | 0.000407014131393417 |
| **BF** | 0.00237605048598787 | 0.000122989861064791 | 0.00987131813842594 | 0.000122868911857295 | 0.00672837511987713 | 0.000267990977032533 |
| **DF** | 0.00235401067661682 | 0.000231380849912071 | 0.0100053813277672 | 0.000231154855181927 | 0.00689687831559475 | 0.000302701913644216 |
| **BC** | 0.00227512354448562 | 0.000465838692709273 | 0.0111043945797906 | 0.00046539299313837 | 0.00759753983710532 | 0.000253904375744097 |
| **CD** | 0.00226904162132366 | 0.000520929502415457 | 0.0112397640076513 | 0.000520420080720753 | 0.00777213702999281 | 0.000289569148222022 |
| **BCF** | 0.00226495985221542 | 0.000294382158550757 | 0.0124925491846874 | 0.000294093039338536 | 0.0117351451259969 | 6.96653501293682E-05 |
| **CDF** | 0.00222293519309333 | 0.00035772597392897 | 0.0126036711834257 | 0.000357372867669789 | 0.0119303465975889 | 7.53498060359215E-05 |
| **D** | 0.00210850764768188 | 0.000414274295669658 | 0.00676622246674496 | 0.000413875877041361 | 0.00256826639562877 | 0.000608178917275148 |
| **EF** | 0.00209114499516206 | 9.55951136427962E-06 | 0.00942914133535863 | 9.5576238781541E-06 | 0.00710428475261598 | 0.000267278326472931 |
| **B** | 0.00209077505758954 | 0.000305199655670175 | 0.00659844078211259 | 0.000304924926314675 | 0.00245188646126821 | 0.000365197674635828 |
| **BCDF** | 0.00207272235316676 | 0.000650742014807366 | 0.013259694831991 | 0.000650025110700575 | 0.015152960182105 | 1.48829495673944E-05 |
| **BDF** | 0.00198686582137387 | 0.000243610502466634 | 0.0115574912958518 | 0.000243344789228769 | 0.0108149387103596 | 6.02714101220126E-05 |
| **BCD** | 0.00196616438884347 | 0.000871540013592 | 0.0125710386537309 | 0.000870577060912887 | 0.0117806753484168 | 5.62216529896832E-05 |
| **CEF** | 0.00189945156591703 | 5.1776610913264E-06 | 0.0120864167982932 | 5.1763656216633E-06 | 0.0121762824532664 | 6.95332552683112E-05 |
| **BD** | 0.0018426523225855 | 0.000365083812397461 | 0.0099007630712008 | 0.000364679826695018 | 0.0067696754129051 | 0.000167308225302824 |
| **BEF** | 0.00182340784621192 | 1.64714490100743E-06 | 0.0110682691508875 | 1.64762451246069E-06 | 0.0110473252151115 | 5.05873727933083E-05 |
| **BCEF** | 0.00181306777966472 | 1.07457093430863E-06 | 0.0128139293879888 | 1.07506132555013E-06 | 0.0154035534551589 | 1.32364833199119E-05 |
| **CE** | 0.00178035309155962 | 2.60910140463736E-06 | 0.0106085595604768 | 2.60779281032329E-06 | 0.00799244195372291 | 0.00025326160791235 |
| **BCDEF** | 0.00168937423224743 | 6.03115724312644E-07 | 0.0132924652833336 | 6.03396320078599E-07 | 0.0180996743057112 | 3.02133422000519E-06 |
| **DEF** | 0.00167340924622634 | 3.16974633489598E-06 | 0.0111709571739063 | 3.16804548871543E-06 | 0.0112384472037267 | 5.65353110472526E-05 |
| **BDEF** | 0.00165760730676499 | 9.095718780067E-07 | 0.0120618453202506 | 9.09817659743428E-07 | 0.0144559153373734 | 1.21692547329166E-05 |
| **CDEF** | 0.0016565897916034 | 2.22417720263646E-06 | 0.0129017493234629 | 2.22296785734854E-06 | 0.0156051663291625 | 1.42906383693811E-05 |
| **BCE** | 0.00157658300666433 | 1.00555709422059E-06 | 0.0120396452911242 | 1.00475853333826E-06 | 0.0120258507158128 | 4.65496962949223E-05 |
| **E** | 0.00154206334705641 | 7.39026314320828E-06 | 0.00604919259180998 | 7.38797568161419E-06 | 0.00271122557889804 | 0.000361641613480834 |
| **BE** | 0.00153915890115875 | 9.25480422030695E-07 | 0.00930856567803832 | 9.2527304048422E-07 | 0.00697635622118008 | 0.000105310087691958 |
| **BCDE** | 0.00148056614567008 | 1.61139226633035E-06 | 0.01288494564619 | 1.60917802004127E-06 | 0.015450722075734 | 1.11671221206048E-05 |
| **CDE** | 0.0014688990370698 | 1.45870071558282E-06 | 0.0121434413739839 | 1.45722458760134E-06 | 0.0122230969743522 | 5.27135017352071E-05 |
| **DE** | 0.00143286574736717 | 1.57316312597442E-06 | 0.00943250568623504 | 1.57201441937875E-06 | 0.00714685696000132 | 0.000142687797535878 |
| **BDE** | 0.00138899463889322 | 8.35438032776333E-07 | 0.0111058253495066 | 8.34675071899989E-07 | 0.0110940424964411 | 3.64621311606214E-05 |
| **NA** | 0 | 0 | 0 | 0 | 0 | 0 |
